# Postnatal intestinal epithelial maturation by LSD1 controls the small intestinal immune cell composition independently from the microbiota

**DOI:** 10.1101/2023.09.08.556818

**Authors:** Alberto Díez-Sánchez, Håvard T. Lindholm, Pia M Vornewald, Jenny Ostrop, Naveen Parmar, Tovah N. Shaw, Mara Martín-Alonso, Menno J. Oudhoff

## Abstract

Postnatal development of the gastrointestinal tract involves the establishment of the commensal microbiota, maturation of the intestinal epithelium, and the acquisition of immune tolerance via a balanced immune cell composition. While studies have uncovered an interplay between the commensal microbiota and immune system development, less is known about the role of the maturing epithelium. Here, we comprehensively show that intestinal-epithelial intrinsic expression of lysine-specific demethylase 1A (LSD1) is necessary for the postnatal maturation of intestinal epithelium as well as maintaining this developed epithelial state in adulthood. Although the stool microbiome was altered in animals with an intestinal-epithelial specific deletion of *Lsd1*, by depleting the microbial component using antibiotics, we found that the cellular state and number of certain immune cell types were dependent on maturation of the epithelium. We found plasma cells, innate lymphoid cells (ILCs), and a specific myeloid population to be depending on epithelial LSD1 expression. We propose that LSD1 controls the expression of epithelial-derived chemokines, such as *Cxcl16*, and this is a mode of action for this epithelial-immune cell interplay. For example, we show that LSD1-mediated epithelial-intrinsic CXCL16 controls the number of local ILC2s but not ILC3s. Together, our findings suggest that the maturing epithelium plays a dominant role in regulating the local immune cell composition, thereby contributing to gut homeostasis.

## INTRODUCTION

Early in life, after birth, the gastrointestinal tract must adapt to a changing environment. There is a nutritional switch from milk to solid foods, while at the same time the microbiota shifts from a few pioneering species in low abundance towards a complex microbial ecosystem with a large load^1^. In addition, during this time, the immune system is educated so it provides tolerance to harmless foreign antigens including commensal microbes^2,3^. Simultaneously, residing in between the microbiota and the immune system, a single layer of intestinal epithelial cells undergoes maturation, which includes the appearance of Paneth cells and progression of other intestinal epithelial cell types into an ‘adult’ state^4,5^. Appropriate postnatal gastrointestinal adaptation is important for our wellbeing. Indeed, necrotizing enterocolitis remains a life-threatening disease for preterm infants^6^, and fetal and early-life antibiotic regimens correlate with developing asthma later in life^7^. In general, gut physiology has been linked to an ever-increasing list of local and systemic diseases ranging from inflammatory bowel disease (IBD), to asthma, and even Alzheimer’s disease^8^.

The small intestinal epithelium dramatically changes in development. Immature villus structures originate at the fetal stage while crypts are formed after birth, finally, during the weaning stage epithelium fully matures which includes the appearance of Paneth cells. This transition also involves a switch from a non-hierarchical at the fetal stage to a hierarchical stem-cell mediated homeostasis of epithelial turnover in adulthood^9^. Changes in WNT and NOTCH signaling or responses to these pathways are important for this early-life switch^9–11^. In addition, modulators of gene expression, especially those controlling persistent gene expression that can putatively alter cellular state and subsequent intestinal epithelial cell composition are involved in this change. For example, the transcriptional repressor BLIMP1/PRDM1 is necessary to prevent early enterocyte maturation and Paneth cell differentiation, and thus *Prdm1*-deficient intestinal epithelium at postnatal day 7 (P7) has accelerated crypt formation and ‘adult’ epithelium with Paneth cells and matured enterocytes^12,13^. In contrast, the Polycomb Repressive Complex 2 (PRC2) member Embryonic Ectoderm Development (EED) likely regulates the mesoderm-to-epithelial transition that occurs prenatally as it represses these genes in adulthood^14^. In addition, the lysine-specific demethylase 1A (KDM1A/LSD1) is required for optimal postnatal Paneth and goblet cell differentiation and *Lsd1*-deficient crypts are neonatal-like^15–17^. However, whether LSD1 also controls other aspects of intestinal epithelial maturation such as enterocyte maturation in villus structures is still unknown.

The immune cell composition in the intestine also undergoes drastic changes in the fetal to adult timeframe. For example, the adult intestine contains macrophage subsets that originated at the perinatal stage and are long-lived, but also subsets that are blood monocyte-derived and recruited after birth that are short-lived and with a high turnover rate^18^. In addition, both fetal- and postnatal-derived type 2 innate lymphoid cells (ILC2s) contribute to the pool of adult gut-resident ILC2s, which have a slow turnover rate during homeostasis^19^. In contrast, Immunoglobulin A (IgA) is absent at birth and IgA-secreting plasma cells rapidly appear in the neonatal stage, after which they continue to be important modulators of the microbiota throughout life^20,21^. Gut immune cell homeostasis is in part coordinated by the epithelium. For example, M cells that sample antigen and cover Peyer’s patches are important for IgA-producing plasma cells^22^, and goblet cells provide antigen during the postnatal stage for regulatory T cell (T_reg_) development^23^ and transfer food antigens throughout life for T_reg_ maintenance^24^. Furthermore, tuft cells, which appear after birth^25^, are the primary source for IL-25 to control ILC2 numbers both during homeostasis and upon infection^26^.

We here provide evidence that LSD1 knockout recapitulates features of neonatal/developing epithelium. This includes the appearance of Paneth cells in crypts, maturation of other secretory cells such as goblet and enteroendocrine cells, as well as maturation of enterocytes in villus structures. Thus, mice with an intestinal-epithelial intrinsic deletion of *Lsd1* (cKO mice) retain a neonatal-like epithelium into adulthood. In addition, deleting *Lsd1* later in life reversed adult matured epithelium towards a neonatal-like state. Postnatal epithelial maturation is instrumental for both the commensal microbial composition and immune cell homeostasis. Macrophage subsets, IgA-expressing plasma cells, and ILCs were remarkably dependent on the maturation state of the intestinal epithelium. Importantly, broad-spectrum antibiotics did not interfere with the several epithelial-immune cell interactions we uncovered, suggesting that epithelial maturation has a dominant role in gut immune cell homeostasis.

## RESULTS

### LSD1 controls differentiation of intestinal epithelial secretory lineages

We have previously shown that lysine-specific demethylase 1A (KDM1A/LSD1) governs the differentiation of intestinal epithelial secretory cells including Paneth and goblet cells^15–17^. In our previous studies, we used a *Villin-*Cre mouse strain that had 10-40% incomplete recombination^27^, which thus led to distinct LSD1**^+^** patches of mature intestinal epithelium^15,17^. As epithelium-derived factors can have a far-ranging effect in modulating its environment, we generated a new mouse line using a different *Villin-*Cre strain^28^ to completely delete *Lsd1* in the intestinal epithelium of both the small intestine (**Fig.1A**) and colon (**Fig. S1A**). Consequently, *Villin*-Cre^+^ *Lsd1*^f/f^ (cKO) mice completely lack Paneth cells throughout the small intestine, as assessed by Lysozyme and Ulex Europaeus Agglutinin 1 (UEA1) staining (**Fig. 1A & 1B**). Furthermore, cKO animals have dramatically reduced numbers of MUC2**^+^** goblet cells in both the small intestine (**Fig. 1A & 1C**) and the colon (**Fig. S1A & S1B**) compared to their wild type littermates. To determine whether other epithelial cell types are also affected by loss of *Lsd1,* we performed bulk RNA-seq from crypts and villi to compare intestinal epithelium from wild type (WT) and cKO animals. Like our previous work (Zwiggelaar *et al.*^15^), we found only modest changes in expression levels of intestinal stem cell markers *Lgr5* and *Olfm4,* as well as the proliferative marker *Mki67,* in crypts (**Fig. 1D & S1C**). In contrast, we found markers for tuft (*Dclk1, Alox5, Il25*), goblet (*Muc2, Clca1*) and enteroendocrine cells (*Chga, Pyy*) to be significantly decreased in villus epithelium (**Fig. 1D & S1C**). This was largely confirmed by performing gene set enrichment analysis (GSEA) of cKO epithelium compared to WT epithelium using cell-specific gene sets from Haber *et al.*^29^ (**Fig. 1E**). Thus, we conclude that cKO mice have crypts without Paneth cells (see also *Lyz1* and *Defa22* in **Fig. 1D & S1C**) and with reduced numbers and/or reduced maturation of all other secretory lineages in the villus, but with intestinal stem cells and proliferative transit-amplifying cells.

**Fig. 1.**
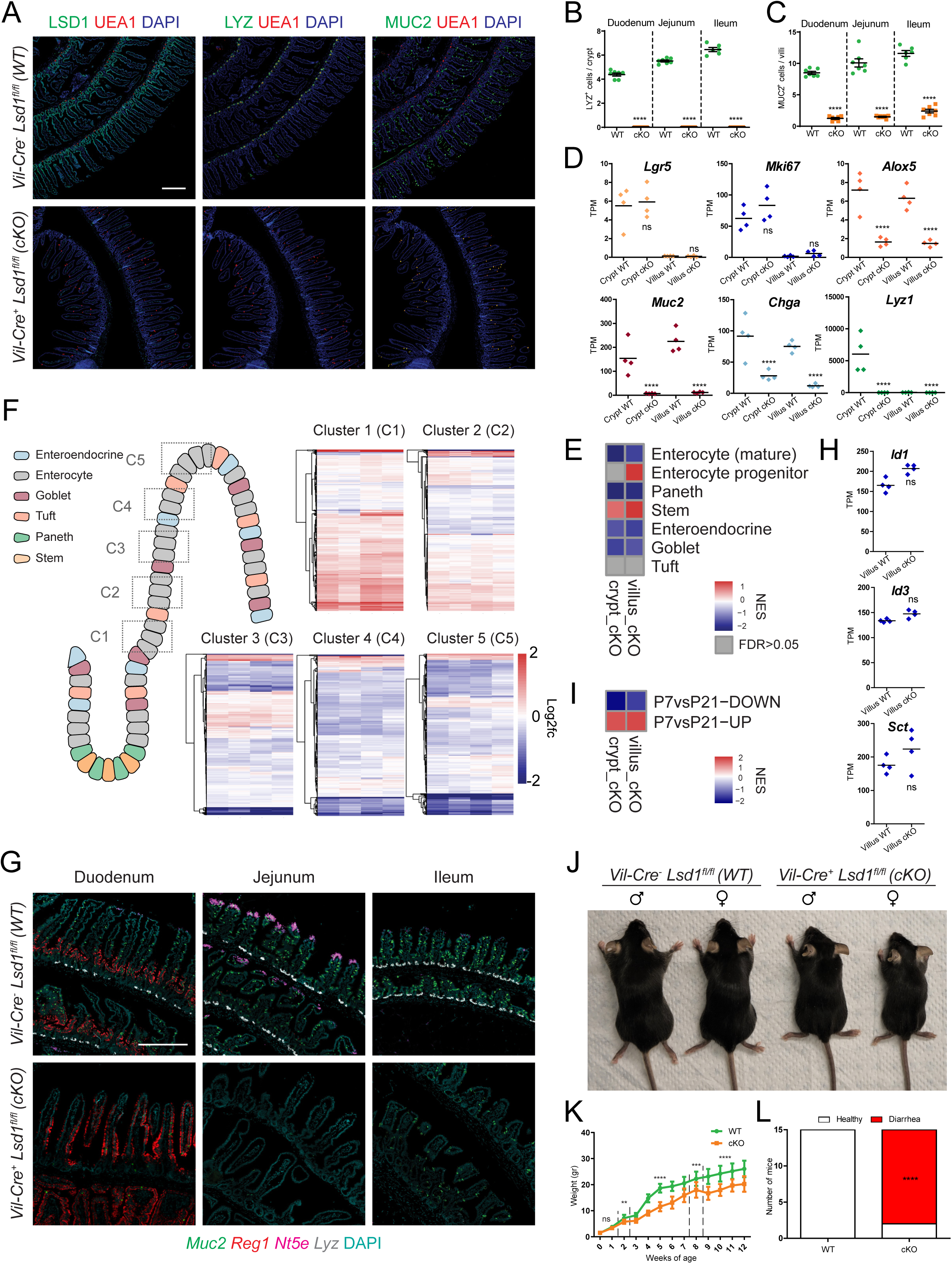
LSD1 is required for the postnatal maturation of the intestinal epithelium. **(A)** Immunofluorescence staining of LSD1, Lysozyme (LYZ), MUC2, and UEA1 of paraffin-embedded duodenal tissue from WT and cKO 2-month-old littermates. Scale bar: 200µm. **(B & C)** Quantification of small-intestinal Paneth (LYZ+) and goblet (MUC2+) cells in WT and cKO mice (derived from images such as shown in Fig. 1A). Data are presented as mean ± SEM; n = 7 mice/genotype from 2 independent experiments, (Two-tailed unpaired t-test). **(D)** Bulk RNA-seq of crypt and villus fractions derived from WT and cKO 2-month-old mice. Individual graphs show Transcripts per Million (TPM). Data are presented as mean & individual data points; n = 4 mice/genotype, (Benjamini–Hochberg adjusted p-value). **(E)** Heatmap showing normalised enrichment scores (NES) from gene set enrichment analysis (GSEA) of cell-type specific gene sets from^29^ (see Table S1) cross-referenced against Fig. 1D bulk RNA-Seq. Crypt cKO compared to crypt WT and villus cKO is compared to villus WT. **(F)** Schematic representation of enterocyte-specific gene expression zonation clusters^30^ along the intestinal villus axis (C1-C5). Heatmaps depict comparison of our villus-derived bulk RNA-seq data against said clusters. Each row is a gene from the gene set and each column represents a biological replicate of villus cKO compared to the mean value of villus WT. Log2fc is capped at -2 and 2. **(G)** Representative fluorescence *in situ* hybridization of cellular markers for goblet (*Muc2*, green), Paneth (*Lyz*, white), villus-base (*Reg1*, red) and villus-tip (*Nt5e*, magenta) enterocytes. Scale bar: 250µm; n = 5 mice/genotype from 2 independent experiments. **(H)** BMP target genes. TPMs derived from Fig. 1D bulk RNA-Seq. Same statistical analysis and data depiction apply. **(I)** Heatmap of NES from GSEA of mouse intestinal epithelium gene expression signatures derived from postnatal day 7 (P7) vs P21 comparisons^15^ (see Table S1) cross-referenced against Fig. 1D bulk RNA-Seq. **(J)** Representative picture of male/female 8-week-old mice. **(K)** Gender aggregated weights are presented as mean ± SEM; n = 12 mice, (6 male & 6 female)/genotype/timepoint from 4 independent experiments, (Individual timepoints assessed by two-tailed unpaired t-test). **(L)** Diarrhea incidence evaluation by stool consistence. n = 15 mice/genotype from 4 independent experiments, (Contingency analysis by Fisher’s exact test). Cell nuclei in all imaging are counterstained with DAPI.

### LSD1 controls enterocyte maturation, independent of BMP signalling

Enterocytes differentiate along the crypt-villus axis, with the least-differentiated enterocytes located at the villus base and the most-differentiated enterocytes near the villus tip, a maturation process mediated by a BMP gradient^30–32^. To assess enterocyte maturation, we plotted Moor *et al.*’s^30^ villus enterocyte-specific zonation gene sets and compared our cKO and WT villus transcriptomes to these maturation signatures. We found an enrichment of ‘clusters 1-2’and overall lower levels of genes associated with the more mature ‘clusters 3-5’ in cKO complete villus epithelium (**Fig. 1F**). In support, by GSEA, we find enrichment of an ‘enterocyte-progenitor’ gene set but repression of a ‘mature-enterocyte’ gene set when comparing cKO and WT villus by bulk RNA-seq (**Fig. 1E**). This suggests that cKO villi are filled with immature enterocytes that would normally only reside at the villus base. To define this in more detail, we performed *in situ* hybridization (ISH) of key cellular marker genes (*Lyz1, Muc2, Reg1, Nt5e, Ada*). Indeed, we find that secretory lineage markers (*Lyz1* and *Muc2*) as well as villus tip enterocyte markers *Nt5e* and *Ada* are reduced in cKO epithelium (**Fig. 1G & S1D**). In addition, we observed a change in distribution for villus-base marker *Reg1*, going from villus base-restricted in WT animals to throughout the duodenal villus structure in cKO littermate animals (**Fig. 1G**). These data may suggest that LSD1 is required for the ability of intestinal epithelium to respond to BMP ligands to mature enterocytes upon the villus axis. However, canonical BMP target genes *Id1* and *Id3* as well as enteroendocrine-specific BMP target gene *Sct*^33^ are equal to or even higher expressed in cKO compared to WT villus epithelium (**Fig. 1H**). Therefore, instead, our data suggests that cKO villus epithelium retains a neonatal-like state into adulthood, rather than the inability to respond to BMP signals. Indeed, by using gene sets originating from transcriptomics comparing intestinal epithelium from postnatal day 7 (P7) *vs.* P21 (from Zwiggelaar *et al.*^15^), we find that genes highly expressed at the P7 stage are enriched in cKO epithelium whereas P21-upregulated genes are repressed (**Fig. 1I**).

In our previous study, we did not find overt abnormalities in *Villin*-Cre *Lsd1*^f/f^ mice (with incomplete deletion), when unchallenged they lived up to at least a year without issues^15^. In contrast, our newly generated cKO animals failed to normally gain weight after weaning (**Fig. 1J & 1K**). In addition, within the first 12 weeks of life, 13 out of 15 cKO animals developed diarrhea (**Fig. 1L**). The timing by which the lack of gaining weight started (weeks 2-4), suggest that cKO animals start to have issues upon switching from milk to solid food, a time window particularly important in intestinal maturation^1^. We envision many factors could cause the lack of weight gain including a lack of properly functioning enterocytes for absorption, impaired enteroendocrine cell function, or a putative altered microbiota. Nevertheless, we conclude that cKO animals retain a neonatal intestinal epithelial cell state into adulthood, which includes absence of Paneth cells, reduced numbers of other secretory cells, and immature enterocytes, yet a clear distinction between crypt and villus, structures that are formed in the first week after birth, remains.

### Mature intestinal epithelium defines and maintains microbial composition

Next, we wanted to test whether maturation of intestinal epithelium affects microbiota establishment and development. We therefore performed 16S ribosomal RNA sequencing of samples isolated from stool and compared WT and cKO litter and cage mate animals at neonatal (P14), before switching from milk to solid food, and adult life stages (6 weeks). By principal coordinate analysis (PCoA) we found the microbiome was indistinguishable at P14, but vastly different at 6 weeks (**Fig. 2A**). Similarly, the microbiota diversity changes (Shannon index) were non-significant when comparing P14 samples but highly significant when comparing adult WT and cKO littermates (**Fig. S2A**). Of note, microbiome in cKO does not retain a neonatal state but develop towards a different composition with notably lesser percentages of *Muribaculaceae* and *Rikenellaceae* and increased percentages of *Bacteroidaceae* and *Sutterellaceae* (**Fig. 2B**).

**Fig. 2.**
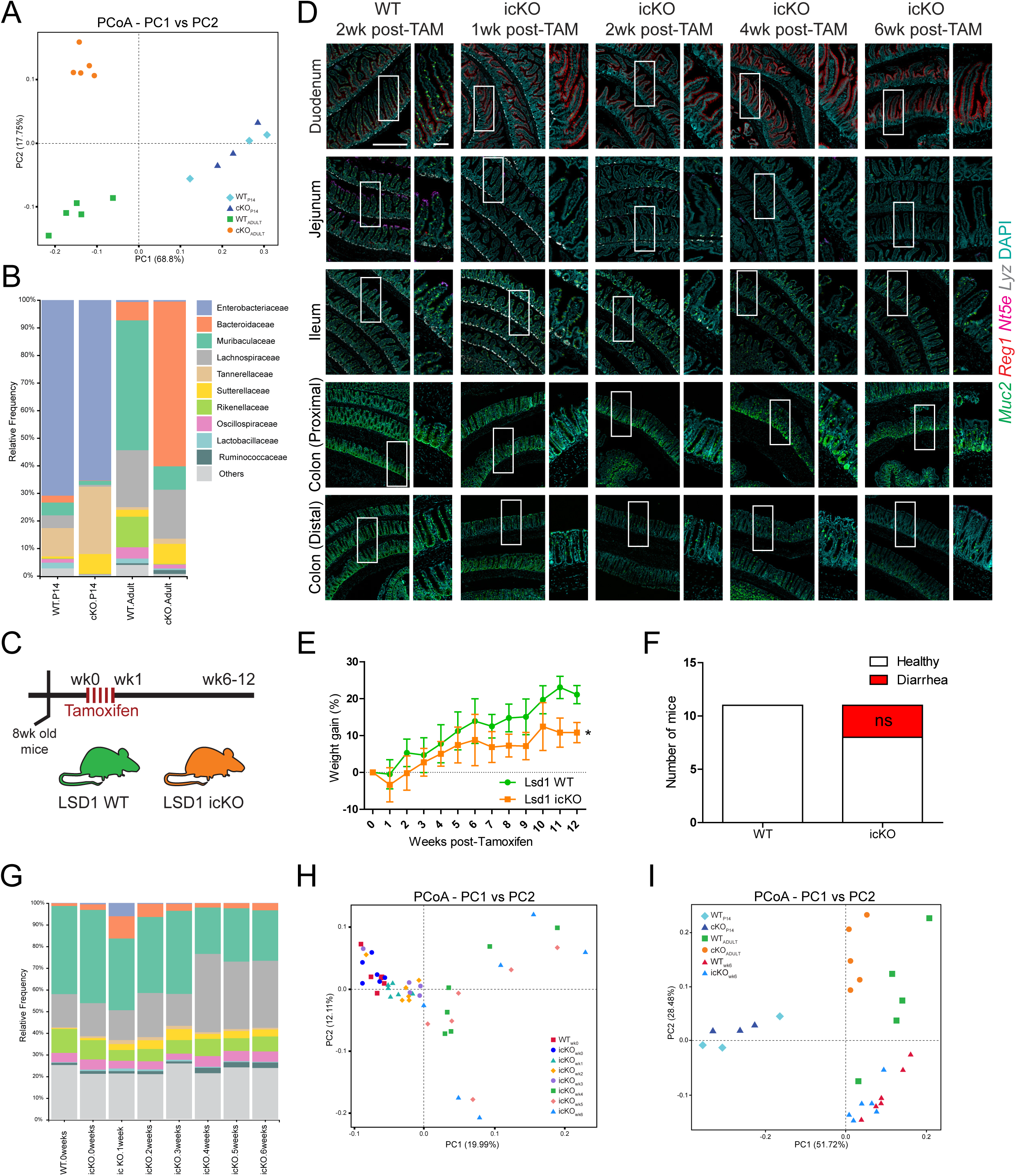
Mature intestinal epithelium defines and maintains microbial composition. **(A)** Principal Coordinate Analysis (PCoA) of stool-derived bacterial 16S rRNA obtained from adult (2-month-old) and P14 mice (WT and cKO); n = 5 adult mice/genotype & n = 3 P14 mice/genotype. **(B)** Average relative frequency of bacterial families found in Fig. 2A stool samples. **(C)** Tamoxifen treatment scheme used to induce complete deletion of *Lsd1* across intestinal epithelium. wk0 = First day of tamoxifen oral gavage, wk1 = 7 days after first tamoxifen oral gavage. Stool and intestinal tissue samples were collected at wk1, wk2, wk4 and wk6 timepoints. **(D)** Representative images of fluorescence *in situ* hybridization of key cellular markers for goblet (*Muc2*, green), Paneth (*Lyz*, white), villus-base (*Reg1*, red) and villus-tip (*Nt5e*, magenta) enterocytes. Scale bar: 500µm, inset scale bar: 100µm; n = 5 mice/genotype from 2 independent experiments. **(E)** Gender aggregated weight assessment of WT and icKO mice post-tamoxifen administration. Weights are presented as mean ± SEM; n = 4 mice, (2 male & 2 female)/genotype/timepoint (Two-way repeated measures ANOVA). **(F)** Diarrhea incidence evaluation by stool consistence. n = 11 mice/genotype from 3 independent experiments, (Contingency analysis by Fisher’s exact test). **(G)** Average relative frequency of bacterial families in stool derived from icKO mice before (icKO.0weeks) and after (icKO.1-6weeks) tamoxifen administration; n = 6 mice/genotype/timepoint **(H)** PCoA representation of Fig. 2G samples. **(I)** Combined PCoA representation of Fig. 2A and Fig. 2H samples.

Maturation of the intestinal epithelium is important for gut microbial composition establishment (**Fig. 2A & 2B**). Next, we wished to test whether matured epithelium is also important for the maintenance of the microbial composition. We therefore generated *Villin*-CreERT2 *Lsd1*^f/f^ (icKO) mice in which we completely deleted *Lsd1* from small intestinal and colon epithelium in adulthood by tamoxifen administration (**Fig. 2C, 2D & S2B**). We found that as early as 2 days after 5 consecutive tamoxifen administration by oral gavage (wk1 timepoint), the intestinal epithelium is completely transformed. We observed a stark reduction in goblet cells, a loss of the mature-enterocyte villus-tip marker *Nt5e* and an expansion of the immature-enterocyte villus-base marker *Reg1* that now covers the whole villus (**Fig. 2D**). Similarly, in the colon, we see a rapid reduction of *Muc2***^+^** goblet cells, especially in the upper half of the crypt (**Fig. 2D**). Although most intestinal epithelium turns over within a week, Paneth cells have an estimated lifetime of 4-6 weeks, and indeed only 4 weeks after *Lsd1* deletion did we observe a complete loss of *Lyz1***^+^** Paneth cells (**Fig. 2D**). Thus, LSD1 is required for Paneth cell differentiation and not their maintenance, unlike other modulators such as ATOH1 where deletion leads to Paneth cell depletion within a week^34^. We hereby establish a model to reverse matured intestinal epithelium towards a neonatal state later in life. As expected, after LSD1 deletion, icKO animals failed to gain weight at the same rate as their WT counterparts, and only 3 out of 11 animals developed diarrhea (**Figs. 2E & 2F**), suggesting that the lack of weight gain in cKO mice is due to a lack of intestinal epithelial maturation and not a consequence of diarrhea.

We collected stool from icKO and WT littermates before and every week after tamoxifen treatment for 6 weeks and performed 16S ribosomal RNA sequencing. Albeit interspecies ecological niche competition and early life colonization being major factors defining gut microbiome composition, upon LSD1 deletion, we recapitulated the same population shifts observed in the cKO, highlighting the importance of the maturation status and/or the presence of a specific adult cellular composition of the intestinal epithelium in modulating the bacterial composition of the gut. Namely an increase in *Bacteroidaceae* and *Sutterellaceae,* and reduced levels of *Muribaculaceae* and *Rikenellaceae* (**Fig. 2G & S2C**). Before tamoxifen administration WT and icKO gut microbiomes were indistinguishable and gradual changes linked to the loss of specific intestinal epithelial cells could be observed at different timepoints (**Fig. 2H**). To name a few recognizable patterns, around the 3 to 4 weeks post-Tamoxifen timepoint, mucus-associated bacteria, such as *Sutterellaceae* and *Bifidobacteriaceae*^35,36^, suffered a sharp decrease coinciding with the loss of most goblet cells across the intestine (**Fig. S2C**). In addition, around the same timepoint, a plethora of opportunistic bacteria (from *Acidaminococcaceae* to *Fusobacteriaceae*) started filling the niche vacuum left by other major players (**Fig. S2C**). Finally, comparing gut microbiomes of WT, cKO and icKO mice we confirm the importance of early life establishment/definition of gut microbial composition, demonstrating how LSD1 deletion in adulthood (icKO), albeit showing the same population shifts, has a much lower impact on changing general bacterial distribution (**Fig. 2I**).

### LSD1 driven epithelial maturation is independent from the microbiota

The microbiota is an established modulator of intestinal epithelial development. For example, microbiota-derived metabolites can alter intestinal epithelial biology^37,38^, and antibiotics-treated animals lack full maturation of intestinal epithelium^39,40^. To test whether the altered microbiota is, in part, responsible for the effect of LSD1 on intestinal epithelium, we treated pregnant mothers and their offspring with a cocktail of antibiotics (ABX, see M&Ms **Fig. 3A**). Like conventionally bred animals (not treated with ABX), we found that cKO animals on ABX did not gain weight properly and most developed diarrhea (**Fig. 3B & 3C**), suggesting that dysbiosis (or changes in the microbiome) did not cause this phenotype.

**Fig. 3.**
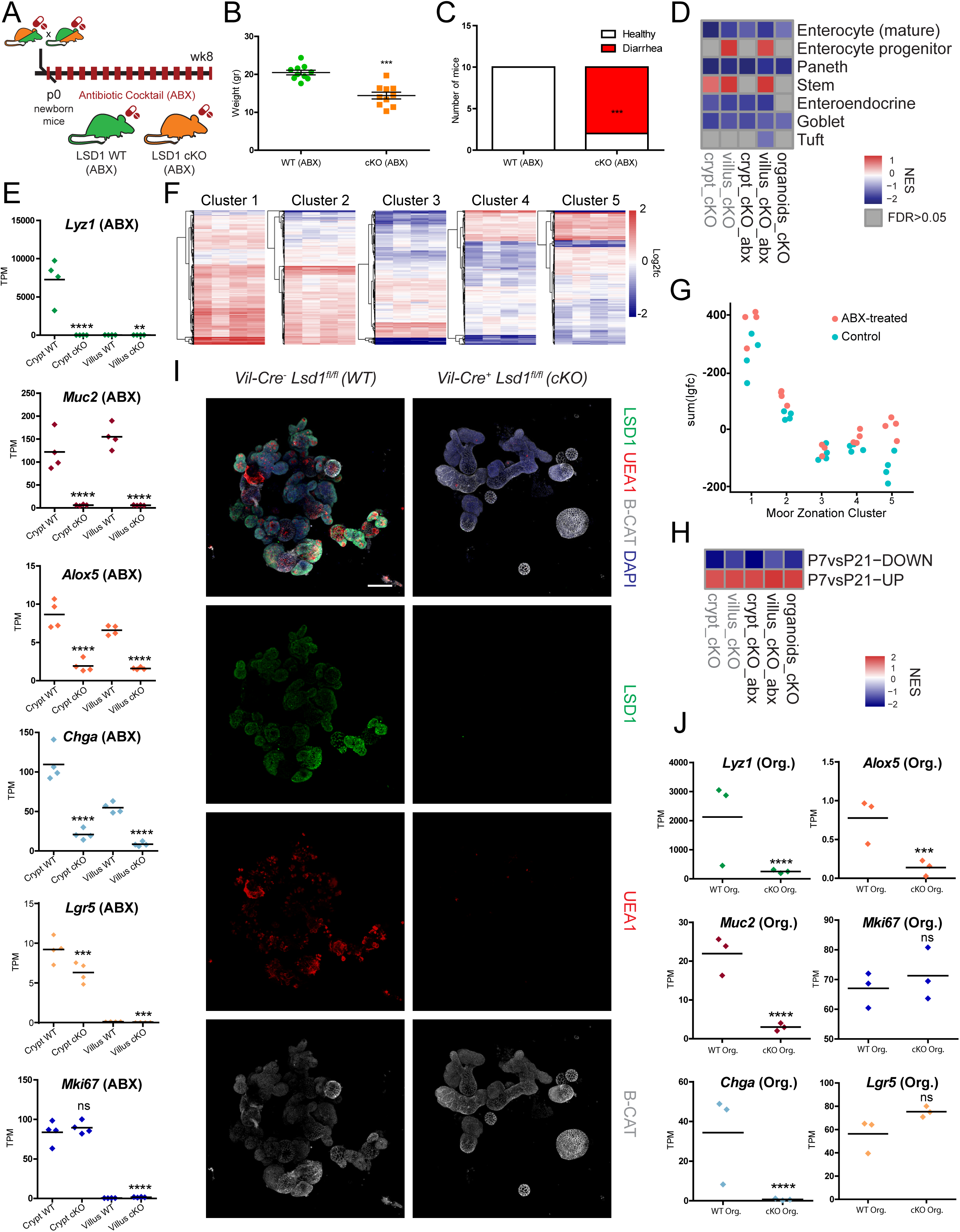
LSD1 driven epithelial maturation is independent from the microbiota. **(A)** Antibiotic cocktail (ABX) treatment scheme. Mating pairs, pregnant mothers and offspring were treated with a cocktail of 5 antibiotics (see M&Ms) during their entire lifetime. **(B)** Gender aggregated weights are presented as mean ± SEM; n = 10 mice, (5 male & 5 female)/genotype from 3 independent experiments, (Two-tailed unpaired t-test). **(C)** Diarrhea incidence evaluation by stool consistence. n = 10 mice/genotype from 3 independent experiments, (Contingency analysis by Fisher’s exact test). **(D)** Heatmap showing normalised enrichment scores (NES) from gene set enrichment analysis (GSEA) of cell-type specific gene sets^29^ (see Table S1) cross-referenced against bulk RNA-Seq of crypts/villi fractions collected from adult WT (ABX) and cKO (ABX) mice. For each comparison, tissue and antibiotic status were matched, i.e. villus LSD1 cKO ABX is compared to villus WT ABX. Bulk RNA-Seq of organoids obtained from untreated WT and cKO mice is included in the comparison. Samples not treated with ABX from Fig.1E are included as reference (grey text). **(E)** Bulk RNA-seq of crypt and villus fractions derived from WT (ABX) and cKO (ABX) 2-month-old mice. Individual graphs show Transcripts per Million (TPM). Data are presented as mean & individual data points; n = 4 mice/genotype, (Benjamini–Hochberg adjusted p-value). **(F)** Heatmaps depict comparison of our villus-derived (ABX) bulk RNA-seq data against enterocyte zonation clusters (see Fig. 1F). Each row is a gene from the gene set and each column represents a biological replicate of LSD1 cKO villus ABX-treated compared to the mean value of WT villus ABX-treated. Log2fc is capped at -2 and 2. **(G)** Dot plot of the summed log2(fold change) in genes from the different zonation clusters from the intestinal epithelium. Each dot represents the sum of a column in Fig. 1F and Fig. 3F. **(H)** Heatmap NES from GSEA of mouse intestinal epithelium gene expression signatures derived from postnatal day 7 (P7) vs P21 comparisons^15^ (see Table S1) cross-referenced against same bulk RNA-Seq data described in Fig. 3D. Untreated samples from Fig. 1I are included as reference (grey text). **(I)** Organoids from untreated WT and cKO mice were cultured for 3 days. LSD1 (green), β-catenin (white) and goblet/Paneth cell (UEA1+) presence was determined by immunofluorescence; n = 3 mice/genotype. **(J)** Bulk RNA-seq of organoids derived from untreated WT and cKO mice. Individual graphs show Transcripts per Million (TPM). Data are presented as mean & individual data points; n = 3 mice/genotype, (Benjamini–Hochberg adjusted p-value).

In addition, by comparing transcriptomes of WT and cKO crypts and villi (derived from ABX mice) we observed a remarkably similar pattern as found in conventionally raised animals: We find complete loss of Paneth cell markers and reduced expression of all other epithelial secretory lineages (**Fig. 3D, 3E & S3A**). Furthermore, we find an enrichment of genes expressed at the villus base (clusters 1-2) and reduced expression of genes associated with the villus tip (clusters 3-5) (**Fig. 3F & 3G**). Using cell-type specific gene sets, we find enrichment of the progenitor enterocyte gene set and repression of a mature enterocyte gene set in ABX-treated cKO animals (**Fig. 3D**), as well as enrichment of P7 and repression of P21 signatures (**Fig. 3H**). Of note, we did observe some differences between conventionally raised and ABX treatment. For example, the stem-cell associated gene *Lgr5* is modestly reduced in ABX treated cKO *vs.* WT crypts whereas it is unchanged in untreated animals (**Figs. 1D & 3E**). In addition, we found upregulation of a number of genes in cKO villus epithelium among the regional gene set clusters 4 and 5 upon ABX treatment (**Fig. 3F & 3G**), whereas almost all those genes were downregulated in conventional animals (**Fig. 1F**). Finally, we isolated organoids from untreated WT and cKO animals and found that also *in vitro*, thus isolated from external factors such as microbiota or immune cells, LSD1 deficient organoids (cKO Orgs.) lacked UEA1-positive Paneth and goblet cells (**Fig. 3I**). Although organoids are considered somewhat immature versions of adult intestinal epithelium, we still observed reduced expression of the mature enterocyte gene set and enrichment of the overall P7 *vs.* P21 gene sets comparing cKO to WT organoids (**Fig. 3D & 3H**) as well as reduced expression of specialized cell markers associated with Paneth, goblet and tuft cells (**Fig. 3J & S3B**).

Together, this body of evidence shows that LSD1 directs the intestinal epithelial maturation process intrinsically, independently of external factors such as the microbiota (ABX-derived data) or the tissue niche (organoid-derived data).

### LSD1-mediated intestinal epithelial maturation does not control systemic immune cell imbalance in spleen or mesenteric lymph nodes

Both the intestinal epithelium and the gut microbiota is dramatically different when comparing adult cKO and WT mice (**Figs. 1-3**). The intestinal epithelium as well as the microbiota are crucial mediators of intestinal and even systemic inflammation. We were therefore surprised that we did not observe any signs of tissue inflammation in cKO animals by gross pathology. For example, splenomegaly and enlarged mesenteric lymph nodes (mLN) are normally associated with a persistent inflammatory response in the gut. However, upon isolation of spleens and mLNs from 6-8 wk old cKO mice and their WT litter mates, we did not observe differences in spleen size (**Fig. 4A & 4B**). In addition, we did not find differences in total cell numbers isolated from mLNs when comparing those from cKO and WT mice **(Fig. 4C)**. Moreover, no overt changes in the immune cell composition (T cells, B cells and antigen-presenting cells) were observed in either spleen-**(Fig. S4A)** or mLN-derived CD45+ cells **(Fig. S4B),** as assessed by flow cytometry. This included CD4**^+^** CD25**^+^** regulatory T (T_reg_) cell numbers **(Fig. S4A & S4B)**.

**Fig. 4.**
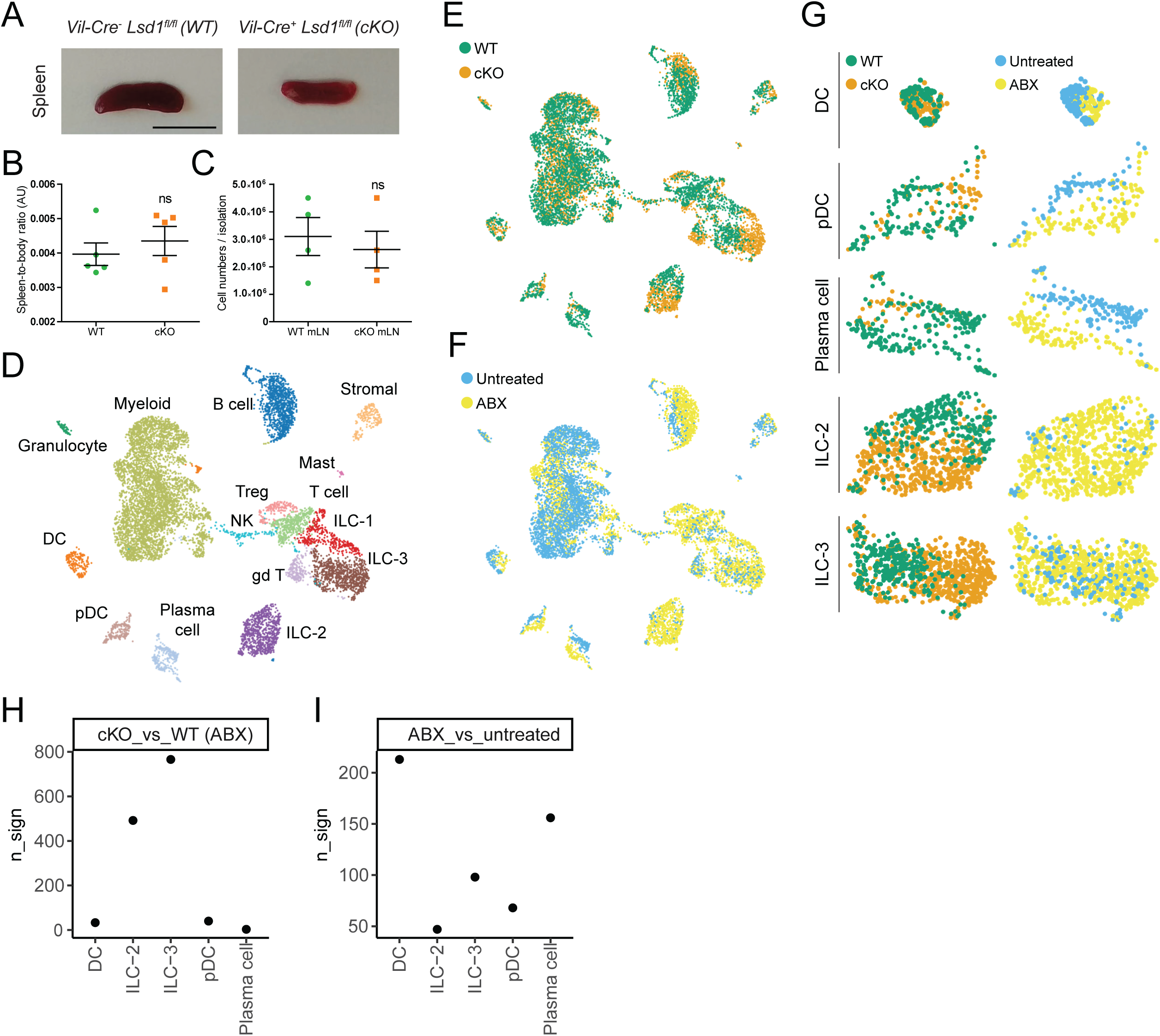
LSD1-mediated intestinal epithelial maturation does not control systemic immune cell imbalance in spleen or mesenteric lymph nodes but directs local immune cell populations. **(A)** Representative picture of spleens derived from 2.5-months-old mice. Scale bar: 1cm; n = 5 mice/genotype. **(B)** Spleen-to-body weight ratio (arbitrary units, AU): n = 5 mice/genotype from 4 independent experiments, (Two-tailed unpaired t-test). **(C)** Number of cells derived from mesenteric lymph nodes (mLN) single cell suspension isolation; n = 4 mice/genotype from 4 independent experiments, (Two-tailed unpaired t-test). **(D)** UMAP (Uniform Manifold Approximation and Projection) plot shows cell types as determined by scRNA-seq from FACS-sorted CD45+ cells derived from the small intestinal lamina propria of ABX and untreated WT and cKO mice. Each immune cell type class is represented as a cluster of points in a unique color; n = 1 mouse/genotype/condition. **(E)** Overlay of genotype (WT or cKO) class over UMAP clustering of all experimental conditions described in Fig.4D. **(F)** Overlay of treatment (untreated or ABX) class over UMAP clustering of all experimental conditions described in Fig.4D. **(G)** Detailed DC, pDC, Plasma cell, ILC2 and ILC3 UMAP clusters derived from Fig. 4E & 4F. **(H)** Number of differentially expressed genes (n_sign, p<0.05) when comparing genotype variable (cKO *vs* WT) in a cell type; n = 1 mouse/genotype (Wilcoxon rank test). **(I)** Number of differentially expressed genes (n_sign, p<0.05) when comparing treatment variable (ABX *vs* untreated, aggregating WT with cKO untreated and WT ABX with cKO ABX samples) in a cell type; n = 1 mouse/genotype/condition (Wilcoxon rank test).

### Intestinal epithelial maturation directs immune cell populations in the lamina propria

To test the role of maturation of intestinal epithelium in the composition of local gut immune cells, we performed single cell RNA-seq (scRNA-seq) on sorted lamina propria CD45**^+^** cells. We were able to distinguish most immune cell types expected to be present (**Fig. 4D & S4C**). As the microbiota is also known to affect local gut immune cell populations and we wanted to dissect the role of intestinal epithelial maturation in the process, we not only performed this experiment in naive cKO and WT animals but also in WT and cKO mice that received ABX (**Fig. 3A & 4D**). Corroborating our finding that cKO mice do not develop overt inflammation, we did not observe drastic changes in the expression of inflammation-associated cytokines (*Ifng, Il1b, Tnf, Il22, Il17a,* and *Il10*) in cells derived from cKO animals compared to those from WT animals (**Fig. S4D**). Next, we grouped all cells either by genotype (*i.e.* WT *vs.* cKO, cells from conventionally bred and ABX treated mice) or by treatment (*i.e.* conventionally bred *vs.* ABX, cells from both WT and cKO mice) (**Fig. 4E & 4F**).

We found that the putative cellular state, as inferred from a change in UMAP clustering, of dendritic cells (DCs), plasmacytoid DCs (pDCs), and IgA-expressing plasma cells is determined by treatment (*i.e.* conventional *vs.* ABX), hence by the presence or absence of microbiota (**Fig. 4G**). In contrast, type 2 and 3 innate lymphoid cells (ILC2s and ILC3s), offer a random distribution when grouping them according to treatment, but display a distinct pattern by genotype, and thus by the maturation state of intestinal epithelium (**Fig. 4G**). In support, we found higher numbers of differentially expressed genes (n_sign) in ILC2s and ILC3s derived from WT and cKO ABX-treated mice as opposed to DCs, pDCs and plasma cells (**Fig. 4H**), while the opposite (except for pDCs) was observed when aggregating WT plus cKO samples and comparing conventional vs. ABX treated mice (**Fig. 4I**).

### LSD1-mediated intestinal epithelial maturation controls intestinal plasma cell numbers and secretory IgA levels

Plasma cells secrete large amounts of IgA to modulate the microbiota and thereby maintain intestinal homeostasis^21^. In our scRNA-seq, we found that there were reduced numbers of plasma cells detected in CD45+ sorted cells from cKO mice compared to WT mice, irrespective of ABX treatment (**Fig. 5A**). Next, we stained small intestinal sections for IgA and confirmed lower number of IgA+ cells in small intestines of cKO compared to WT (**Fig. 5B**). Similarly, IgA levels in small intestinal content were significantly lower in those derived from cKO animals compared to WT animals, irrespective of treatment (**Fig. 5C**). Overall, ABX treatment led to reduced numbers of IgA+ cells and lower levels of IgA in intestinal content (**Fig. 5B & 5C**). In part, plasma cells originate in Peyer’s patches where specialized epithelial M cells transfer antigen from the lumen, and mice lacking M cells have reduced IgA levels in a similar manner to what is observed in our cKO animals^22^. Although we were not able to stain for these cells, we deem it likely that M cell maturation, alike all other epithelial lineages, is much impaired in cKO animals. Indeed, in our bulk RNA-seq data, we found lower levels of the M-cell marker gene *Spib* in cKO epithelium compared to WT epithelium (**Fig. S5A**). Expectedly, *Spib* is a gene that only starts to be expressed in intestinal epithelium between postnatal day 7 and 21 (**Fig. S5B**). In future work, we hope to address M cell differentiation, antigen uptake, and plasma cell development in more detail.

**Fig. 5.**
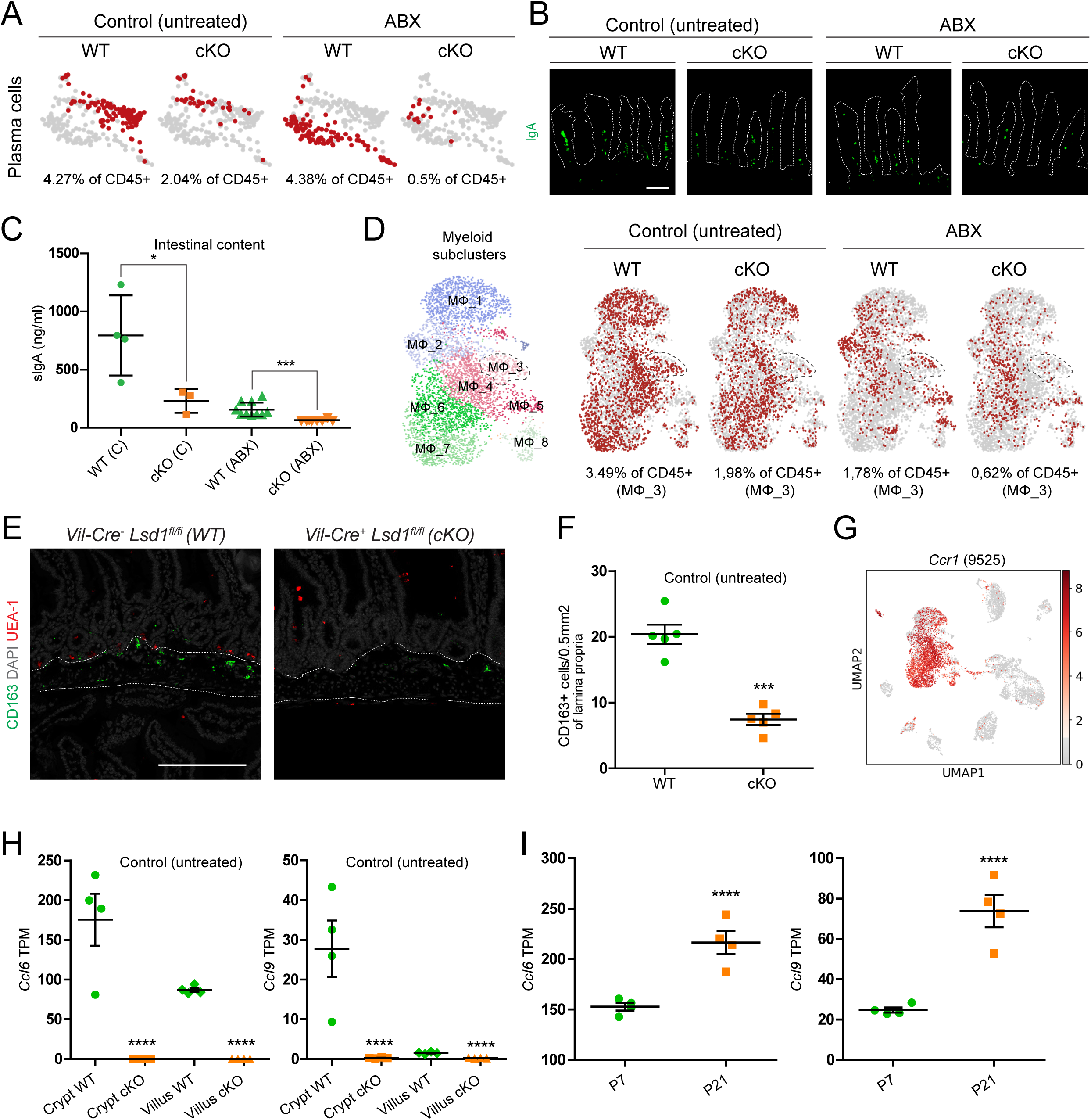
LSD1-mediated intestinal epithelial maturation controls intestinal plasma cell and macrophage homeostasis. **(A)** UMAP clustering of lamina propria plasma cells across all experimental conditions. Red dots correspond to the number of plasma cells detected under each condition. Grey dots represent all detected plasma cells across conditions. **(B)** Immunofluorescence staining of paraffin-embedded mouse duodenal tissue depicting IgA-producing plasma cells in green. Scale bar: 100µm. Villus structure is delimited by a discontinuous white line; n = 5 mice/genotype/condition from 2 independent experiments **(C)** Secreted IgA (sIgA) protein concentration as determined by ELISA on small intestinal content of 2-month-old WT and cKO mice (untreated and ABX-treated). Data are presented as mean ± SEM; n = 4 mice/genotype (untreated) & n = 10 mice/genotype (ABX) from 3 independent experiments, (Two-tailed unpaired t-test). **(D)** UMAP subclustering of lamina propria myeloid population in 8 different populations (MΦ_1-8) across all experimental conditions. MΦ_3 is highlighted across conditions by a discontinuous line (% of CD45 corresponds to this subcluster). **(E)** Paraffin section of mouse duodenum submucosal region. CD163+ macrophages (green) are detected by immunofluorescence, UEA1+ cells (red) correspond to Paneth and/or goblet cells. Nuclei counterstained with DAPI (grey). Scale bar: 200µm; n = 5 mice/genotype from 2 independent experiments (untreated). CD163+ cell density in the intestinal submucosal region; n = 5 mice/genotype from 2 independent experiments (untreated). **(G)** UMAP of lamina propria CD45+-derived cells showing *Ccr1* gene expression across all experimental conditions. Number in between parentheses represents the number of sequenced cells that passed quality control, (9525) corresponds to all four conditions merged under one UMAP. Grey to red heatmap scale shows log(CPM+1). **(H)** Bulk RNA-seq of crypt and villus fractions derived from WT and cKO untreated 2-month-old mice. Individual graphs show Transcripts per Million (TPM). Data are presented as mean ± SEM; n = 4 mice/genotype, (Benjamini–Hochberg adjusted p-value). **(I)** Bulk RNA-seq of intestinal epithelium derived from WT mice at postnatal day 7 (P7) vs P21^15^ . Individual graphs show Transcripts per Million (TPM). Data are presented as mean ± SEM; n = 4 mice/timepoint, (Benjamini–Hochberg adjusted p-value).

### LSD1-mediated intestinal epithelial maturation is required for a putative CCL6/CCL9-CCR1 axis in macrophage establishment or recruitment

Initially, in our scRNA-seq UMAP clustering we identified one large myeloid population (**Fig. 4D**). Subsequent re-clustering of this population in isolation led to eight myeloid subclusters (MΦ_1-8) (**Fig. 5D**). Upon assessing individual panels, we found that ABX treatment drastically reduces the percentage of total myeloid cells, as is commonly found by others^41,42^. In contrast, myeloid population 2 (MΦ_2) appeared upon ABX treatment (**Fig. 5D**). We were unable to assess whether this is due to a shift in an existing population, or an appearance and recruitment of a separate one. Next, we focused on changes in cKO compared to WT derived cells. In addition to overall lower percentage of myeloid cells in cKO, we found that myeloid population 3 (MΦ_3) was nearly absent (**Fig. 5D**). This population expressed specific markers such as *F13a1* and *Cd163* (**Fig. S5C**). In a recent preprint, CD163 was identified as a marker for a long-lived macrophage subset residing underneath intestinal crypts^43^. Indeed, upon staining with an anti-CD163 antibody, we confirmed this specific location and reduced numbers in the small intestine of cKO mice compared to those from WT mice (**Fig. 5E & 5F**).

Chemokine receptors are important factors in macrophage gut homeostasis, indeed the CCL2-CCR2 axis is crucial for monocyte recruitment from blood which subsequently differentiate into macrophages in the tissue^44,45^. Within intestinal tissue, similarly to other tissues, CCR2^+^ and CCR2^-^ Mϕ populations co-exist, identifying recently differentiated Mϕ and long-lived Mϕ, respectively^46^. Our population of interest, MΦ_3, included a mixture of *Ccr2* expressing and non-expressing cells (**Fig. S5C**). Notably, however, most myeloid cells expressed *Ccr1*, including MΦ_3 (**Fig. 5G & S5C**). In search of changes in corresponding chemokine receptor-ligand expression of the intestinal epithelium, we found that *Ccl6* and *Ccl9,* which encode CCR1 receptor ligands CCL6 and CCL9 respectively, were nearly completely absent in cKO epithelium, irrespective of ABX treatment (**Fig. 5H & S5D**). In line with the submucosal localization of CD163^+^ macrophages, both *Ccl6* and *Ccl9* show higher expression in adjacent crypt fractions compared to villus fractions (**Fig. 5H & S5D**). Indeed, both these chemokines are expressed in goblet and Paneth cells as deducted from published scRNA-seq data from Haber *et al.*^29^ (**Fig. S5E**), and for CCL6 this crypt location was found previously^47^. Importantly, both *Ccl6* and *Ccl9* are markers that are drastically induced at P21 compared to P7 (**Fig. 5I**), and thus part of the early-life intestinal maturation that is governed by LSD1 and may contribute to establishment of CD163^+^ macrophages within this niche.

### LSD1-mediated intestinal epithelial maturation is required for the establishment and maintenance of ILC2s and ILC3s in a microbiota-independent manner *via* a putative CXCL16-CXCR6 axis

We found that the cellular state, as inferred from UMAP clustering and differential gene expression, of small intestinal ILC2s and ILC3s is dependent on genotype (**Fig. 4G & 4H**). Overall, gut resident ILCs arrive, expand, and/or mature early in life and are tuned by the presence of the microbiota^19,48^. Therefore, our next step was to assess ILC2 and ILC3 numbers in adult tissue. We found higher relative ILC2 and ILC3 numbers in the CD45+ population by scRNA-seq when comparing WT with cKO derived cells (**Fig. S6A**). Next, we quantified cells using fluorescent immunostaining of small intestinal sections. ILC2s were detected and quantified using CD3-RORγt-GATA3+ as markers and ILC3s were defined as CD3-RORγt+ entities. In support of the scRNA-seq data, both these cell types were detected in increased quantities in cKO mice compared to WT mice, both in the presence (untreated controls) and absence (ABX) of microbiota (**Fig. 6A, 6B, 6C, S6B, S6C & S6D**). Finally, increased numbers of ILC3s were also confirmed by surface staining using flow cytometry (**Fig. 6D**).

**Fig. 6.**
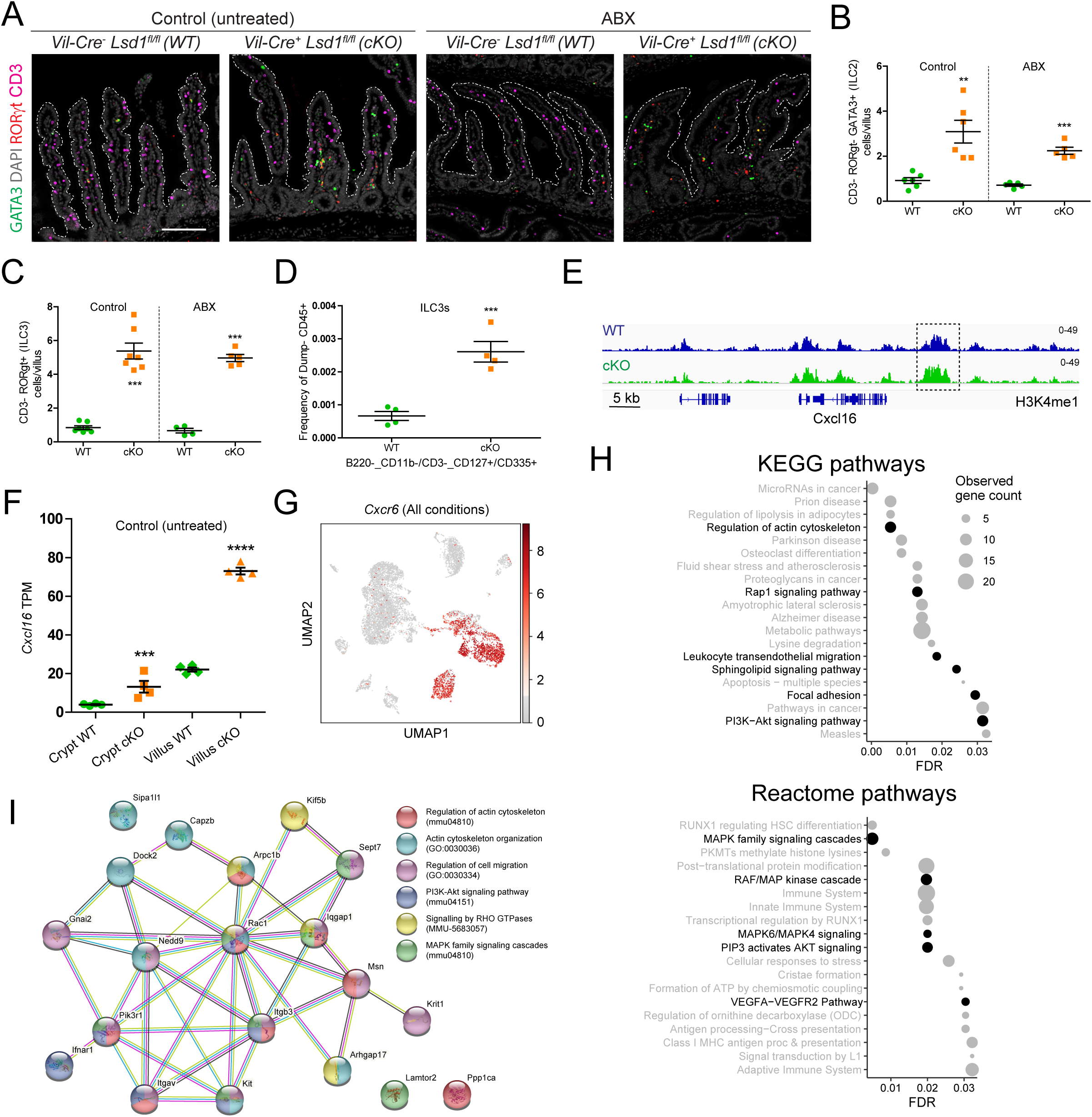
LSD1-mediated intestinal epithelial maturation is required for the establishment and maintenance of ILC2s and ILC3s in a microbiota-independent manner. **(A)** Type 2 and 3 Innate Lymphoid Cells (ILC2s & ILC3s) detection by immunofluorescence in mouse duodenum of WT and cKO mice (untreated and ABX-treated). ILC2 cells are defined as CD3^-^ RORγt^-^ GATA3^+^ while ILC3s are CD3^-^ RORγt^+^. Nuclei are counterstained with DAPI (grey). Villus structure is delimited by a discontinuous white line; Scale bar: 200µm. See Fig. S6B for split channels. **(B)** Quantification of ILC2s in the lamina propria of duodenum from untreated and ABX-treated mice (derived from Fig. 6A images). Data are presented as mean ± SEM; n = 6 mice/genotype (untreated) and 5 mice/genotype (ABX-treated) from 2 independent experiments, (Two-tailed unpaired t-test). **(C)** Quantification of ILC3s in the duodenal lamina propria of untreated and ABX-treated mice (derived from Fig. 6A images). Data are presented as mean ± SEM; n = 7 mice/genotype (untreated) and 5 mice/genotype (ABX-treated) from 2 independent experiments, (Two-tailed unpaired t-test). **(D)** Flow cytometry data derived from small intestinal lamina propria CD45+ cells showing ILC3 (Ly6g^-^/B220^-^ CD11b^-^/CD3^-^ CD127^+^[IL-7Rα^+^]/CD335^+^[NKp46^+^]) frequency. Each data point represents an individual mouse, independent experiments were carried out for each WT and cKO pair, (Two-tailed unpaired t-test). Full gating hierarchy is shown in (Fig. S4E). **(E)** Representative Integrative Genomics Viewer (IGV) tracks of H3K4me1 levels near the *Cxcl16* locus (n = 2). Box indicates a significantly increased peak. **(F)** Bulk RNA-seq of crypt and villus fractions derived from WT and cKO untreated 2-month-old mice. Individual graphs show Transcripts per Million (TPM) counts. Data are presented as mean ± SEM; n = 4 mice/genotype, (Benjamini–Hochberg adjusted p-value). **(G)** Lamina propria CD45+-derived UMAP showing *Cxcr6* gene expression across all experimental conditions. Grey to red heatmap scale shows log(CPM+1). **(H)** Enrichment analysis of the 140 genes commonly upregulated in ILC2s and ILC3s derived from ABX-treated cKO mice and performed with the STRING online tool. Top 20 pathway hits with the lowest false discovery rate (FDR) are displayed. Canonical pathways activated by the CXCL16-CXCR6 axis are highlighted in black. **(I)** StringDB-derived interactome showing a 20 gene subset (of first and second order Rac GTPase interactors) from the 140 gene list described in Fig.6H and S6G. Participation of each gene in a particular biological process is shown as a color overlay (GO, MMU and mmu prefix codes correspond to Gene Ontology: Biological Processes, Reactome pathways and KEGG pathways databases respectively). For interaction color coding see M&Ms.

The intestinal epithelium is known to be able to locally control ILCs by expressing specific factors. For example, ILC2s respond to epithelial-derived IL-25, IL-33 and TSLP, while IL-15 and IL-18 can promote the number of ILC3s. Although we observed higher levels of *Il18*, we found equal or lower expression of all other putative epithelial factors that could control these cell types (*Il25, Il33, Tslp, Il15*) when comparing cKO epithelium with that from WT (**Fig. S6E**). Given that most of the factors know to influence ILC2s and ILC3s abundance were unchanged or reduced, we were prompted to assess other genes encoding cytokines and chemokines capable of interacting with ILC2s and ILC3s and controlled by LSD1, in an unbiased manner. In a previous study, we performed H3K4me1 ChIP-seq on small intestinal crypts from WT and cKO animals (note that these cKO animals did not have complete *Lsd1* deletion^15^). Here, we assessed this data set in search for significantly increased H3K4me1 levels in or near genes encoding for a cytokine or chemokine and combined this with receptor expression patterns in ILC2s and ILC3s from our scRNA-seq. We found increased levels of H3K4me1 near the gene that encodes for the chemokine CXCL16 (*Cxcl16*) (**Fig. 6E**), suggesting a direct effect of lack of demethylation by LSD1. In support, *Cxcl16* was expressed at significantly higher levels (∼4-fold) in the intestinal epithelium of cKO mice compared to that of WT mice, particularly in villus epithelium and irrespective of ABX treatment (**Fig. 6F & S6F**). CXCL16 is a ligand for CXCR6, and, indeed, *Cxcr16* was expressed in our populations of interest (ILC2s and ILC3s) among other cell types (**Fig. 6G**).

Finally, we assessed what genes were differentially expressed in ILC2s and ILC3s by genotype using the scRNA-seq data set. We identified 140 genes that were commonly upregulated in both ILC2s and ILC3s derived from the lamina propria of cKO mice compared to WT mice (**Fig. S6G**). This overlap in upregulated genes in these two different cell types suggests that this could be due to a common activation, such as through the CXCL16-CXCR6 axis. Indeed, we found that many of the commonly upregulated genes are associated with processes canonically activated downstream of CXCL16-CXCR6 signalling such as actin cytoskeletal reorganization and subsequent chemotaxis, PI3K-Akt, MAPK cascade, and RHO GTPase signalling^49^ (**Fig. 6H**). To conclude, a detailed assessment of these 140 genes unearthed a network of Rho/Rac GTPases, actin cytoskeleton regulators and integrins commonly found in cell adhesion and migration processes (**Fig. 6I**). Altogether, these results point to enhanced recruitment of ILC2s and ILC3s to the lamina propria of cKO mice, a process putatively driven by aberrant production of CXCL16 by cKO epithelium.

### ILC2s and ILC3s rapidly accumulate upon intestinal-epithelial *Lsd1* deletion in adult mice

Taking advantage of our inducible cKO system (icKO, **Fig. 2C**), we tested if reverting matured epithelium into a neonatal-like state in adult animals would affect ILC2s and ILC3s in the small intestine. Strikingly, as early as 2 days after 5 daily consecutive tamoxifen administrations by oral gavage (wk1 timepoint, see Fig. 2), we observed an increase in ILC2s and ILC3s in the lamina propria of icKO mice as compared to WT animals that received the same treatment (**Fig. 7A, 7B & S7A**). The increase in ILC2s and ILC3s in icKO animals was persistent throughout the 6-week experiment (**Fig. 7A, 7B & S7A**). Of note, these fast changes in cell numbers take place at a time when the microbiota of icKO mice is practically unchanged as compared to that of a WT, which further strengthens the microbiota-independent effect (see icKOwk1, **Fig. 2H**).

**Fig. 7.**
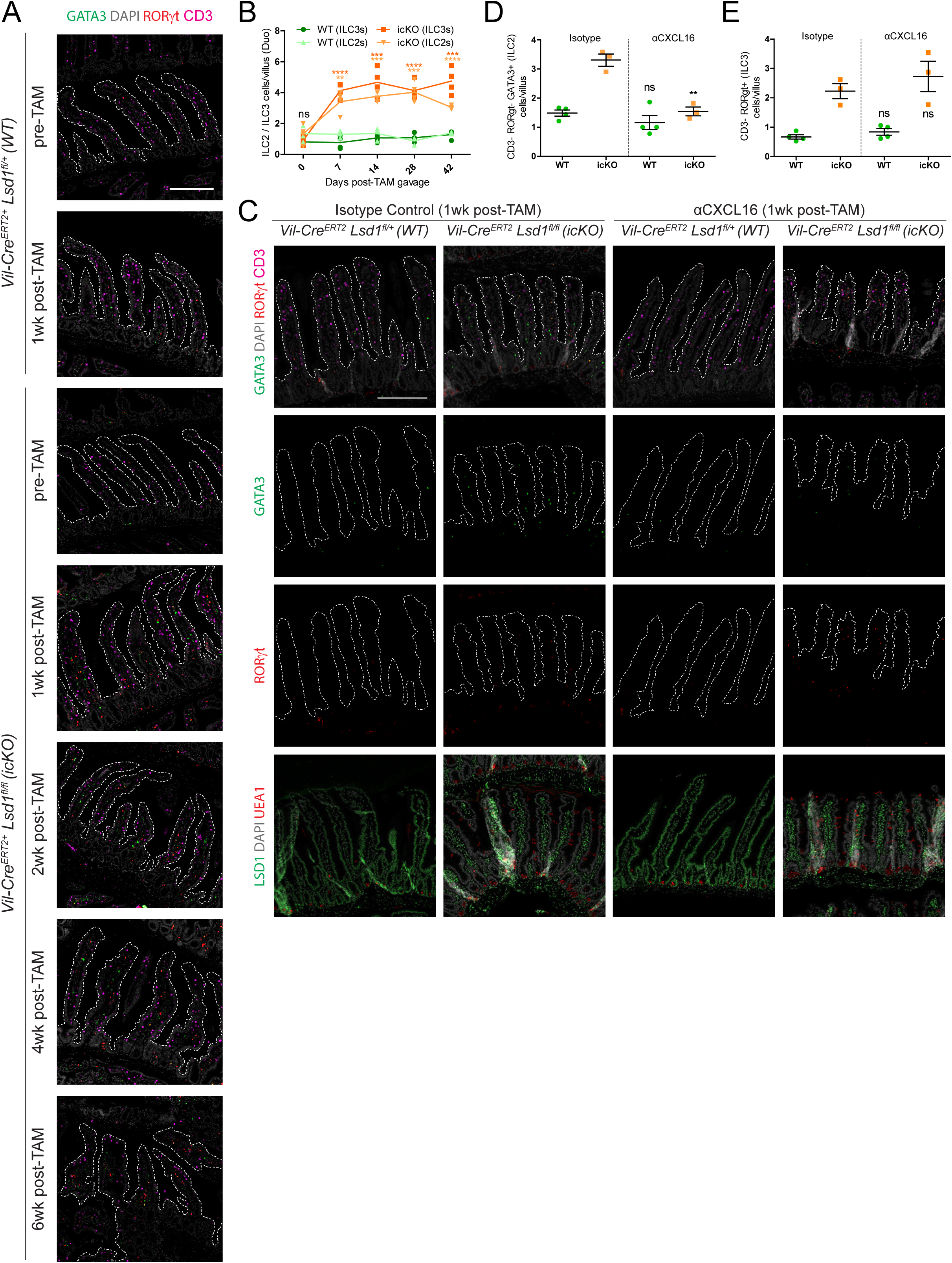
Rapid induction of CXCL16-dependent ILC2s and CXCL16-independent ILC3s upon epithelial-intrinsic *Lsd1* deletion in adult mice. **(A)** ILC2s and ILC3s detected by immunofluorescence in mouse duodenum of WT and icKO mice; before tamoxifen treatment (pre-TAM) and after 1 to 6 weeks of initial tamoxifen dosage (post-TAM 1-6wk). ILC2 cells are defined as CD3^-^ RORγt^-^ GATA3^+^ while ILC3s are CD3^-^ RORγt^+^. Nuclei are counterstained with DAPI (grey). Villus structure is delimited by a discontinuous white line; Scale bar: 200µm. See Fig. S7A for split channels. **(B)** Quantification of ILC2s and ILC3s in the lamina propria of duodenums from pre- and post-TAM treated mice (derived from Fig. 7A images). Data are presented as individual data points in a time series; n = 4 mice/genotype/timepoint (Individual timepoints/cell type assessed by two-tailed unpaired t-test). **(C)** ILC2s and ILC3s detected by immunofluorescence in mouse duodenum of 1wk post-TAM WT (subject to Isotype control or αCXCL16 treatment, as shown in Fig. S7B). ILC2 cells are defined as CD3^-^ RORγt^-^ GATA3^+^ while ILC3s are CD3^-^ RORγt^+^. Nuclei are counterstained with DAPI (grey). Villus structure is delimited by a discontinuous white line; Scale bar: 200µm. **(D)** Quantification of ILC2s in the lamina propria of duodenum from Isotype control and αCXCL16-treated mice (derived from Fig. 7C images). Data are presented as mean ± SEM; n = 4 mice/genotype (Isotype control) and 3 mice/genotype (αCXCL16-treated), (Two-tailed unpaired t-test). **(E)** Quantification of ILC3s in the lamina propria of duodenum from Isotype control and αCXCL16-treated mice (derived from Fig. 7C images). Data are presented as mean ± SEM; n = 4 mice/genotype (Isotype control) and 3 mice/genotype (αCXCL16-treated), (Two-tailed unpaired t-test).

### Anti-CXCL16 treatment blocks ILC2 but not ILC3 accumulation upon intestinal-epithelial *Lsd1* deletion in adult mice

To test if the hypothesized CXCL16-CXCR6 axis was the main contributing factor governing the increase of ILC2s and ILC3s in the lamina propria of cKO and icKO mice we devised a signalling blockade experiment (**Fig. S7B**) using a monoclonal antibody targeting CXCL16 (αCXCL16). While ILC2 numbers were significantly affected by CXCL16 blockade (**Fig.7C and 7D**), ILC3s still increased after blocking CXCL16 (**Fig. 7C and 7E**) in icKO mice. Of note, we did not observe any changes in WT animals receiving the antibody treatment, suggesting that short-term ILC2/3 maintenance in adulthood is not mediated by CXCL16, which is in line with work by others^50^. Our study thus indicates that LSD1-mediated repression of *Cxcl16* is necessary to limit ILC2 numbers. In contrast, ILC3 abundance might be controlled by other concomitant factors.

## DISCUSSION

We here show that LSD1 is required for the postnatal maturation of epithelium that lines the small and large intestine (**Figs. 1-3**). Postnatal intestinal epithelial maturation comprises the appearance of Paneth cells and expansion and/or maturation of all other cell lineages including goblet cells and enterocytes. This role for LSD1 seems tissue specific, as it opposes its role in skin epithelium where inhibition of LSD1 led to the promotion of differentiation^51^. The microbiota can modulate intestinal epithelial cell biology, including Paneth cell numbers, for example through the production of metabolites altering histone de-acetylase activity^38,52^. However, we find that LSD1 has an intrinsic and dominant role, as ABX treatment *in vivo* or culturing epithelium *in vitro* in organoids did not overtly affect the impaired intestinal epithelial maturation in the absence of LSD1 (**Fig. 3**). Using transcriptomic analysis, we find that adult *Lsd1*-deficient small intestinal epithelium is enriched for genes highly expressed at P7 and repressed by P21, the weaning stage (**Fig. 1I & 3H**). We do observe normal crypt formation though, which is initiated in the first week of life, suggesting that up to P7 LSD1 does not play a major role in intestinal epithelial development. In contrast, the polycomb repressive complex 2 (PRC2) member EED (Embryonic ectoderm development) is necessary to repress genes normally expressed at the embryonic stage, thus prior to the formation of gut epithelium in development^14^. Indeed, deletion of EED in intestinal epithelium renders adult mice moribund within 2 weeks after the first tamoxifen administration, including the loss of proliferative crypt structures^14^. Thus, where EED is responsible for the mesoderm-to-epithelial transition, LSD1 is needed for the postnatal maturation transition. Together, epigenetic modifiers are strongly involved at different stages in intestinal epithelial development and epigenetic enzymatic activity can be microbiota-dependent and independent.

The intestinal epithelium is an important modulator of the commensal microbial composition. Goblet cells produce mucins to form a protective layer, Paneth cells secrete antimicrobials such as lysozyme and defensins, and enterocytes express *Reg3b* and *Reg3g* in a microbiota-dependent manner^29,53^. Expectedly, the microbiome of adult cKO animals was different from its WT littermates (**Fig. 3A & 3B**). In addition, deletion of *Lsd1* in adulthood (icKO mice) led to changes in the microbiome in a similar manner, albeit in a much more modest fashion (**Fig. 3G & 3I**). As the change in icKO animals was increasing in a week-by-week manner, we could not conclude whether specific cell types, such as Paneth cells that were only fully gone by four weeks after deletion, are dominant in their role controlling the microbiome.

The intestinal epithelium is intertwined with local immune cells to maintain gut homeostasis. Using scRNA-seq, we tested what immune cell types were microbiota-dependent and, by ABX treatment, what aspects of such immune cell composition were defined by LSD1 controlled epithelial maturation (**Figs. 4-7**). We found that IgA-producing plasma cell numbers were controlled both by epithelial maturation status and the microbiota (**Fig. 5A-C**). Goblet and M cells are involved the transfer of antigen from the lumen^22–24^ and this is likely to be affected in cKO mice. In addition, although macrophage cellular state, as inferred by UMAP clustering, was mostly unaffected, we determined that the establishment or maintenance of a CD163+ macrophage population was dependent on intestinal epithelial maturation. Potentially, via the crypt-enriched expression of chemokines *Ccl6* and *Ccl9* by goblet and Paneth cells (**Fig. 5H, S5D & S5E**), which are ligands for macrophage-expressed CCR1 (**Fig. 5G**). Furthermore, both the cellular state as well as the number of ILC2s and ILC3s in the lamina propria is controlled by epithelial-expressed *Lsd1*. We propose that LSD1 normally represses *Cxcl16* as we found heightened levels of H3K4me1 near its locus (**Fig. 6E**) and strongly increased *Cxcl16* expression in cKO epithelium compared to WT epithelium (**Fig. 6F & S6F**). ILC2s and ILC3s express *Cxcr6*, the CXCL16 receptor (**Fig. 6G**), and thus we initially hypothesized that heightened *Cxcl16* levels could cause both ILC2 and ILC3 recruitment or expansion. Indeed, we find that a gene set upregulated in both ABX cKO-derived ILC2s and ILC3s is associated with pathways activated by a CXCL16-CXCR6 axis (**Fig. 6H & 6I**). In support, mice lacking *Cxcr6* have reduced numbers of ILC3s in homeostasis^54^ and the CXCL16-CXCR6 axis has been shown to govern migration in ILC2s^55–57^. In our study, the changes in ILC numbers occured quite rapidly as deletion of *Lsd1* in adult epithelium led to an increase of ILC2s and ILC3s within a week (**Fig. 7A, 7B & S7A**). Of note, CXCL16 expression is normally associated with dendritic cells^54,58^, but as epithelium is by far the most prevalent cell type in the gut, increased expression could lead to high protein levels.

Given the results of our *in vivo* blocking experiments we present the CXCL16-CXCR6 axis as a key driver behind the ILC2 accumulation upon *Lsd1* deletion in the intestinal epithelium. In contrast, ILC3 accumulation was unaffected in αCXCL16 treated mice (Fig. 7). While most epithelial-derived factors normally associated with controlling ILC2 and ILC3 cell numbers were not strikingly increased and therefore unlikely contributors (**Fig. S6E**), *Il18*, a cytokine described to drive ILC3 proliferation^59^, was significantly upregulated in cKO intestinal epithelium (**Fig. S6E**). This would potentially allow ILC3s to circumvent the effects of CXCL16 neutralization via the IL18-IL18R pathway.

The results of this study confirm that LSD1 is not only necessary for the correct maturation of the intestinal epithelium but also for the maintenance of this mature status. In addition, we show a new gut-microbiota-independent axis of communication between the intestinal epithelium and the immune cell compartment, a process directly influenced by the maturation status of said epithelium. Finally, although this study provides the backbone for direct communication between the intestinal epithelium and several immune cell types, further intervention-type work on ligand-receptor interactions blockade is needed to characterize these in more depth.

## MATERIAL AND METHODS

### Animal experiments

#### Ethics statement

Mice were maintained at the Comparative Medicine Core Facility (CoMed) at NTNU (Norges Teknisk Naturvitenskaplige Universitet) in accordance with the Norwegian Guidelines for Animal Research. Antibiotic cocktail and tamoxifen induced deletion experiments were assessed and approved by the Norwegian Food Safety Authority (FOTS ID 21275).

#### Mouse strains

*Villin*-Cre (Jackson Laboratories, Strain #: 021504, RRID:IMSR_JAX:021504), *Villin*-CreERT2^27^ (kind gift from Dr. Robine) and Lsd1f/f^60^ (kind gift from Stuart Orkin) mice were housed in CoMed. Villin-Cre Lsd1f/f mice were housed under specific-pathogen free (SPF) conditions and Villin-CreERT2 Lsd1f/f mice were maintained in the minimal disease unit at CoMed.

#### Mice euthanasia method

Euthanasia of rodents was done via carbon dioxide (CO2) inhalation. In order to cause minimal distress and obtain rapid unconsciousness mice were euthanized in their home cage A fill rate of 30-70% of the chamber volume per minute with 100% CO2, added to the existing air in the chamber, was used to achieve a balanced gas mixture

#### Antibiotic Cocktail Treatment (ABX)

Parental mice were treated with antibiotic cocktail (ABX) in drinking water 2 weeks prior to mating and subsequent offspring kept being administered ABX for the duration of the experiment. ABX was freshly prepared weekly, kept at 4°C and changed every 2-3 days. Antibiotic cocktail (ABX): Ampicillin (A0166-5g, Sigma-Aldrich) (1g/liter), Amoxicillin-Clavulanate (SMB00607-1G, Sigma-Aldrich) (0,25g/liter), neomycin (Colivet vet., VetPharma AS) (1g/liter), gentamicin (G1397-100 ml, Sigma-Aldrich) (1g/ liter), and vancomycin (V2002, Sigma-Aldrich) (0,5g/liter) in drinking water. Sucralose (sweetener) was added at 0,1g/liter to make the cocktail palatable.

#### Tamoxifen administration

Tamoxifen (Sigma-Aldrich, T5648-1G) was dissolved in 50ml of Corn Oil (Sigma-Aldrich, C8267-500ML) at a 20 mg/ml concentration and stored at 4°C. To induce *Lsd1* recombination, 8–12-week-old mice were administered daily via oral gavage with 0.1ml of Tamoxifen for a duration of 5 days.

#### CXCR6-CXCL16 signalling blockade experiment (αCXCL16)

To induce *Lsd1* recombination, 8–12-week-old mice were administered daily via oral gavage with 0.1ml of Tamoxifen for a duration of 5 days. Monoclonal rat IgG2A CXCL16 neutralizing antibody (R&D Systems, Clone #142417, MAB503) or monoclonal Rat IgG2A isotype control (R&D Systems, Clone #54447, MAB006) was administered via intraperitoneal injection at a dose of 100μg/mouse (in 200 μl of sterile PBS) for 4 alternating days, starting the day before the first tamoxifen oral gavage. Antibodies were freshly reconstituted in sterile PBS on the day of the injection. See Fig. S7B for treatment scheme.

### Intestinal epithelium isolation

#### Crypt and villi isolation

Briefly, we isolated 10 cm from the duodenal section of mice intestine. The duodenum was washed with ice-cold PBS, open longitudinally and cleaned of mucus with a coverslip. Next, villi were scraped off using a glass coverslip and collected with ice-cold PBS. Collected villi were filtered through a 70µm cell strainer and cleaned with 20ml of ice-cold PBS. By inverting the filter villi were recovered using PBS and centrifuged at 300 × g for 5 min. Villi fractions were snap-freezed in liquid nitrogen for subsequent RNA extraction. Afterwards, we followed standard protocol for crypts isolation as described^61^. Remaining duodenum tissue was cut into 2–4 mm small pieces and transferred to a 50 mL tube. These fragments were washed with ice cold PBS by up and down with a 10mL pipette coated with FBS. This step was repeated 10 times until supernatant turned fully clear. Next, these fragments were incubated with 30 mL ice-cold crypt isolation buffer (2mM EDTA in PBS) with gentle rotation in the cold room. After 30 minutes, fragments were settled down and supernatant was discarded. After this, fragments were vigorously shaken with 20ml of PBS to release the villus and crypts. This step was repeated until most of crypts were released and passed through the 70-μm cell strainer and collected into FBS coated sterile 50ml falcon tube. The filtered fractions contain crypts and were spun down at 300 × g for 5 min and pellet was resuspended in 10 mL of ice-cold basal culture media. This fraction was washed with ice-cold basal culture media to remove single cells at 200 × g for 2 mins. Next, pellets were either snap-freezed for RNA extraction or resuspended and counted under light microscope for seeding as an organoid culture.

#### Organoid cultures

Organoids were generated by seeding ca. 250–500 small intestinal crypts in a 40 μl droplet of cold Matrigel (#734-1101, Corning) into the middle of a pre-warmed 24-well plate. Matrigel was solidified by incubation at 37°C for 5–15 min and 500 μl of basal culture medium (ENR) added. Basal culture medium (“ENR”) consisted of Advanced DMEM F12 (Gibco) supplemented with 1x Penicillin-Streptomycin (Sigma-Aldrich), 10 mM HEPES, 2 mM Glutamax, 1x B-27 supplement, 1x N2 supplement, (all Gibco) 500 mM N-Acetylcysteine (A7250, Sigma-Aldrich), 50 ng/ml recombinant EGF (PMG8041, Thermo Fisher Scientific), 10% conditioned medium from a cell line producing Noggin (kind gift from Hans Clevers), and 20% conditioned medium from a cell line producing R-Spondin-1 (kind gift from Calvin Kuo). ENR culture medium was replaced every 2–3 days. Organoids were passaged at 1:3–1:4 ratio by disruption with rigorous pipetting almost to single cells. Organoid fragments were centrifuged at 300 × g, resuspended in 40 μl cold Matrigel per well, and plated on pre-warmed 24-well plates.

### Fluorescence confocal microscopy staining, imaging and quantification

#### Immunofluorescence staining of paraffin-embedded tissue and organoids

After sacrificing the mice, 10cm of representative sections of the intestinal tract were dissected, washed with ice-cold PBS and rolled into “swiss rolls” as previously described^62^. Intestinal tissue was fixed with 4% formaldehyde for 24h at RT (Room Temperature). After fixation, tissues were paraffin-embedded, cut in 4-μm sections and placed in slides. Tissue slides were then deparaffinized and rehydrated, followed by antigen retrieval (Citrate buffer pH-6,0 for all stainings). Sections were blocked in blocking/permeabilization buffer (0,2% Triton X-100, 1% BSA and 2% normal goat serum (NGS) in 1X PBS) for 1 hr at RT in a humidified chamber. Tissues were then incubated overnight at 4°C in staining buffer (0,1% Triton X-100, 0.5% BSA, 0,05% Tween-20 and 1% NGS in PBS) with diluted primary antibodies: (LSD1, 1:200, Cell Signaling Technology, Cat. No. 2184S), (MUC2, 1:200, Santa Cruz Biotechnology, sc-15334), (LYZ, 1:200, Agilent DAKO, A0099), (β-CAT, 1:200, BD Biosciences, 610154), (CD3, 1:200, Novusbio, NB600-144155), (RORγt, 1:200, ThermoFisher, 12-6988-82), (GATA3, 1:200, Abcam, ab199428), (CD163, 1:200, Abcam, ab182422), (CD68, 1:200, BioRad, MCA1957T) and (IgA, 1:200, Novusbio, NB7503). Next day, slides were washed with 0.2% Tween-20 in PBS. After 3 washing steps, tissues were incubated with secondary antibodies coupled to fluorochromes, UEA-1 (1:500, Vector Laboratories, RL-1062) and counterstained with Hoechst 33342 (1:1000, MERCK, H6024) for 1 hr at RT. After incubation, slides were washed 3 times with 0.2% Tween-20 in PBS and then mounted in Fluoromount G (Invitrogen, 00-4958-02). For signal amplification of CD163 and multiplexing with primary antibodies from the same species (Rabbit, CD3 and GATA3) the Tyramide SuperBoost™ (ThermoFisher, B40922) kit was used. For organoid staining we followed an already published protocol^17^.

#### RNAScope (*In Situ* RNA analysis)

To perform the 4-plex *in situ* hybridisation on pretreated formalin-fixed paraffin-embedded tissues both RNAScope Multiplex Fluorescent Kit v2 (ACD Biotechne, Cat. No. 323100) and 4-Plex Ancillary Kit (ACD Biotechne, Cat. No. 323120) kits were used in combination with Opal reagents from Akoya Biosciences (Opal 520 – Cat. No. FP1487001KT, Opal 570 – Cat. No. FP1488001KT, Opal 620 – Cat. No. FP1495001KT, Opal 690 – Cat. No. FP1497001KT) for detection of fluorescent signals. The following probes were used for target detection: (Mm-Ada-O1, Cat. No. 832851), (Mm-Nt5e, Cat. No. 437951), (Mm-Lyz-C2, Cat. No. 415131-C2), (Mm-Muc2-C3, Cat. No. 315451-C3) and (Mm-Reg1-C4, Cat. No. 511571-C4). Tissue processing and staining workflow was carried out according to RNAScope 4-plex Ancillary Kit for Multiplex Fluorescent Reagent Kit v2 technical note.

#### Confocal Microscopy Imaging

Images were obtained on a Zeiss LSM880 Airyscan using two air objectives: Plan-Apochromat 10x/0.45NA/2mm WD (420640-9900) and a Plan-Apochromat 20x/0.8NA/0.55mm WD (420650-9902). Imaging was performed at the Cellular and Molecular Imaging Core Facility (CMIC), Norwegian University of Science and Technology (NTNU).

#### Image Acquisition, Processing Software and Quantification

Images were acquired using Zen 2.3 Black and grey levels/maximum intensity projections were adjusted in Zen 2.3 Blue prior to .tiff export. Cell counts were manually quantified in an unbiased manner, blind to sample genotype, using sample numbers as identifiers. Depicted scale bars apply to all images within the same panel.

### Lamina propria, spleen and mesenteric lymph node leukocyte isolation

After sacrificing 2.5 months old mice, lamina propria leukocyte isolation (from the entire small intestine length) was done according to the referenced method^63^ with a few modifications, namely: HBSS was substituted with DPBS (w/o CaCl2 & MgCl2) and the tissue digestion was performed on 15ml of prewarmed (37°C) RPMI 1640 containing: 10% FCS + Pen/Strep (10mM) + Glutamine (2mM) + Collagenase D 7.5mg + Dispase 45mg and DNAse 7.5mg + 100ul COLLAGENASE VIII (62.5 CDU/ml). For spleen isolations, the spleen was minced using scissors and then ground using the flat end of a syringe in 5ml of RPMI on a 100 ml culture dish. The resulting cell suspension was passed through a 40um cell strainer into a 50ml tube. Cells were spun down, supernatant removed, cell pellet resuspended in Red Blood Cell Lysis Buffer (eBioscience, Cat. No. 00-4333-57) and incubated for 2 minutes at RT. Afterwards, lysis reaction was diluted with 45ml of RPMI, cells spun down and resuspended in PBS + 10% FBS for surface staining. Mesenteric lymph nodes were harvested, with care to remove any remaining fat, and stored in cold PBS with 10% FCS. Harvested lymph nodes were passed through 70 μm cell strainer to obtain a single cell suspension. Cell-numbers and viability were assessed using Trypan blue in an EVE automated cell counter (VWR).

### Flow cytometry

Single cells were stained with Zombie Aqua (Biolegend, 1:1,000 in PBS) for 15 min at room temperature (RT) for live-dead exclusion. Samples were incubated with two different panels (St1 and St2), consisting of antibody conjugates against: [St1: CD335-BV421, CD3-BV605, CD127-BV711, CD8-BV785, CD25-AF488, TCRgd-PerCp-Cy5.5, TCRb-PE, CD4-APC, CD45-APC-Fire, Dump (CD326, CD19, CD11b, Ly6g, Ter119)-PE-Cy7] and [St2: CD335-BV421, CD3-BV605, CD127-BV711, CD25-BV785, MHCII-AF488, CD11c-PerCp-Cy5.5, B220-PE, CD11b-AF647, CD45-APC-Fire, Dump(CD326, Ly6g, Ter119)-PE-Cy7] [all Biolegend, 1:200 in PBS + 2% fetal calf serum (FCS)] for 20 min at 4°C. All the samples were analyzed using a BD LSR II flow cytometer (BD Biosciences) equipped with 405, 488, 561, 647 nm laser lines. Single fluorochrome stainings of cells and compensation particles (BD CompBead, Becton Dickinson) were included in each experiment. For analysis, FlowJo software v10.6.2 and GNU R/Bioconductor v3.6.3/v3.10 packages flowCore, CytoML/flowWorkspace, ggcyto, flowViz were used (Van *et al*., 2018).

### Nucleic acid extractions, sequencing and analysis

#### RNA extraction

RNA was isolated from organoids using the RNeasy Plus Micro Kit (Qiagen, Cat. No. 74034) and from intestinal epithelium (crypts and villi) using the RNeasy Plus Mini Kit (Qiagen, Cat. No. 74134) in combination with QIAshredder columns (Qiagen, Cat. No. 79656). All samples were snap frozen in liquid nitrogen on isolation day and purified simultaneously later.

#### gDNA extraction from mouse stool

Genomic DNA extraction from frozen stool (snap frozen in liquid N2) samples was done using the QIAamp Fast DNA Stool Mini Kit (Qiagen, Cat. No. 51604) according to protocol. Lysis temperature was adjusted to 95℃.

#### Bulk RNA-Seq sequencing

Library preparation and sequencing were performed by Novogene UK using the NEB Next Ultra RNA Library Prep Kit (New England BioLabs, Cat. No. E7530L). Samples were sequenced using 150-bp paired-end reads using NovaSeq 6000 (Illumina).

#### Bulk RNA-Seq analysis

Reads were aligned to the Mus musculus genome build mm10 using the STAR aligner^64^. The count of reads that aligned to each exon region of a gene in GENCODE annotation M18 of the mouse genome^65^ was counted using featureCounts^66^. Genes with a total count less than 10 across all samples were filtered out. Differential expression analysis was done with the R package DESeq2, and volcano plots were plotted with the R package EnhancedVolcano^67^. PCA analysis was performed with the scikit-learn package with the function sklearn.decomposition.PCA^68^. GSEA analysis was run with the log2(fold change) calculated by DESeq2 as weights, 10,000 permutations, and otherwise default settings using the R package clusterProfiler^69^. GO term analysis was run using the function enrichGO in clusterProfiler. The R packages pheatmap and eulerr were used to make heatmaps and venndiagrams, respectively.

#### scRNA-Seq sequencing and analysis

scRNA-seq was performed using 10X genomics Chromium Next GEM Single Cell 3′ GEM, Library & Gel Bead Kit v3.1 and sequenced using two Illumina NS500HO flowcells with a 75 cycle kit. Reads were aligned to the genome and reads per gene was counted using the cellranger count function (v3.1.0) with the cellranger reference mm10 3.0.0^70^. Samples were then combined with cellranger aggr. Scanpy (v1.5.1)^71^ was used to quality filter the raw data with the following filters: min_genes_per_cell = 400, min_cells_per_gene = 3, max_counts_per_cell = 40000, max_percent_mt_per_cell = 20. Raw reads were converted to counts per million and log-transformed. Cell neighbours were computed with 10 neighbours and 40 PCA dimensions. Leiden algorithm was used with a resolution of 1 to split clusters and UMAP to display them. CIPR with the ImmGen dataset from mouse was used to initially identify the cell clusters^72^, and these were confirmed with cell type markers. Scanpy was used to make plots of cell clusters.

#### StringDB gene enrichment

Version 11.5 of STRING Database was used for gene enrichment and network graph display. Interaction sources to build up the network include: Textmining, experiments, databases, co-epxression, neighbourhood, gene fusion and co-occurrence. Only experimentally determined (magenta) and known interactions (cyan) from curated databases are depicted along interactions derived from textmining (green) and co-expression (black).

#### Microbiome sequencing

16S rDNA Amplicon Metagenomic Sequencing of bacterial gDNA derived from mouse stool was outsourced to Novogene UK. Amplicon library of the V4 region was obtained using the 515F (5’-GTGCCAGCMGCCGCGGTAA-3’) and 806R (5’-GGACTACHVGGGTWTCTAAT -3’) primer pairs connected with barcodes. The PCR products with proper size were selected by 2% agarose gel electrophoresis. Same amount of PCR products from each sample was pooled, end-repaired, A-tailed and further ligated with Illumina adapters. Paired-end sequencing (PE250) was performed on the obtained libraries based on the Illumina NovaSeq sequencing platform.

#### Microbial 16S Amplicon analysis

Amplicon sequence analysis was outsourced to Novogene UK. Paired-end reads was assigned to samples based on their unique barcodes and truncated by cutting off the barcode and primer sequences. After the barcode sequence and primer sequence were truncated, FLASH (v1.2.11, http://ccb.jhu.edu/software/FLASH/)^73^ was used to merge the reads to get raw tags. Then, fastp software was used to do quality control of raw tags, and high-quality clean tags were obtained. Finally, Vsearch software was used to blast clean tags to the database to detect the chimera and remove them, to obtain the final effective tags. For the effective tags, the DADA2 or deblur module in QIIME2 software was used to do denoise (DADA2 were used by default), and the sequences with less than 5 abundance were filtered out to obtain the final ASVs (Amplicon Sequence Variables) and feature table. Then, the Classify-sklearn moduler in QIIME2 software was used to compare ASVs with the database and to obtain the species annotation of each ASV. QIIME2 software was used to calculate alpha diversity indices including observed_ otus, shannon, simpson, chao1, goods_ coverage, dominance and pielou_e indices. The rarefaction curve and species accumulation boxplot were drawn. If there was grouping, the differences between groups of alpha diversity would be analyzed by default. For Beta Diversity analysis, the UniFrac distance was calculated by QIIME2 software, and the dimensionality reduction maps of PCA, PCoA and NMDS were drawn by R software. Among them, Ade4 and ggplot2 packages in R software were used to display PCA and PCoA. Then, Adonis and Anosim functions in QIIME2 software were used to analyze the significance of community structure differences among groups. Finally, LEfSe or R software was used to perform the species analysis of significant differences between groups. LEfSe analysis was performed by LEfSe software, and LDA score threshold was set to 4 by default. In MetaStat analysis, R software was used to test the differences between the two groups at the level of phylum, class, order, family, genus and species, and P value was obtained. The species with P value less than 0.05 were selected as the significant differences between the two groups; in T-test, R software was also used to analyze the significant differences of species at each taxonomic level.

### Statistical Analysis

Unless stated otherwise in the methods section, statistical analysis and representation was carried out using Graphpad Prism V8.0. Statistical significance was evaluated by the appropriate methods stated in the figure legends or in methodology section. Means were represented as ±SEM unless stated otherwise. Differences were considered as statistically significant at *P<0.05, **P<0.01, ***P<0.001, and ****P<0.0001.

### Data availability

All sequencing data was uploaded via Annotare to the ArrayExpress repository at BioStudies EMBL-EBI (www.ebi.ac.uk/arrayexpress) under the following accession numbers. For bulk RNA-Seq-derived data: E-MTAB-10498 (Crypt and villi small intestinal epithelium from untreated WT and cKO mice), E-MTAB-10499 (Crypt and villi small intestinal epithelium from ABX-treated WT and cKO mice) and E-MTAB-10500 (Passage number 4, day 4 organoids derived from the duodenum of untreated WT and cKO mice). For scRNA-Seq-derived data: E-MTAB-10492. For 16S rDNA Amplicon Metagenomic Sequencing of bacterial gDNA derived from mouse stool: E-MTAB-12489 (Adult vs Neonate) and E-MTAB-12626 (Timepoints before and after *Lsd1* tamoxifen induced recombination in the intestinal epithelium).

## Supporting information

Genesets

## ACKNOWLEDGEMENTS

We thank Drs. Stuart Orkin and Sylvie Robine for kindly sharing mouse strains. We thank Unni Nonstad and Liv Ryan for assistance with cell sorting. We thank the imaging (CMIC) and animal care (CoMed) core facilities that assisted in this work (NTNU). The single-cell RNA-seq was done by the Genomics Core Facility at NTNU, which receives funding from the Faculty of Medicine and Health Sciences and Central Norway Regional Health Authority. Funding of this work was provided by the Research Council of Norway (Centre of Excellence grant 223255/F50, and ‘Young Research Talent’ 274760 to MJO and 326209 to MMA) and the Norwegian Cancer Society (182767 to MJO). TNS was funded by the BBSRC (BB/S01103X/2).

## Author contributions

Conceptualization of the study: ADS and MJO; Performed experiments: ADS, PMV, MMA, JO and NP; Analysis and interpretation of data: ADS, MMA, HTL, TNS and JO; Bioinformatic analysis: HTL; Made figures: ADS, MMA, HTL and PMV; Supervision: MJO; Wrote first draft: ADS, MMA and MJO. Contributed to final draft: ADS, HTL, PMV, NP, JO, TNS, MMA and MJO.

### Competing interests

Correspondence and request for materials should be addressed to MJO

## Notes

### Competing Interest Statement

The authors have declared no competing interest.

## REFERENCES

1. Dogra, S. K. et al. Nurturing the Early Life Gut Microbiome and Immune Maturation for Long Term Health. Microorganisms 9, (2021).

2. Zheng, D., Liwinski, T. & Elinav, E. Interaction between microbiota and immunity in health and disease. Cell Research 2020 30:6 30, 492–506 (2020).

3. Peterson, L. W. & Artis, D. Intestinal epithelial cells: regulators of barrier function and immune homeostasis. Nature Reviews Immunology 2014 14:3 14, 141–153 (2014).

4. Elmentaite, R. et al. Cells of the human intestinal tract mapped across space and time. Nature 2021 597:7875 597, 250–255 (2021).

5. Fawkner-Corbett, D. et al. Spatiotemporal analysis of human intestinal development at single-cell resolution. Cell 184, 810–826.e23 (2021).

6. Hunter, C. J., Upperman, J. S., Ford, H. R. & Camerini, V. Understanding the susceptibility of the premature infant to necrotizing enterocolitis (NEC). Pediatr Res 63, 117–123 (2008).

7. Örtqvist, A. K. et al. Antibiotics in fetal and early life and subsequent childhood asthma: nationwide population based study with sibling analysis. BMJ 349, (2014).

8. Blumberg, R. & Powrie, F. Microbiota, disease, and back to health: a metastable journey. Sci Transl Med 4, (2012).

9. Guiu, J. et al. Tracing the origin of adult intestinal stem cells. Nature 570, 107–111 (2019).

10. Fordham, R. P. et al. Transplantation of expanded fetal intestinal progenitors contributes to colon regeneration after injury. Cell Stem Cell 13, 734–744 (2013).

11. Mustata, R. C. et al. Identification of Lgr5-independent spheroid-generating progenitors of the mouse fetal intestinal epithelium. Cell Rep 5, 421–432 (2013).

12. Muncan, V. et al. Blimp1 regulates the transition of neonatal to adult intestinal epithelium. Nat Commun 2, (2011).

13. Harper, J., Mould, A., Andrews, R. M., Bikoff, E. K. & Robertson, E. J. The transcriptional repressor Blimp1/Prdm1 regulates postnatal reprogramming of intestinal enterocytes. Proc Natl Acad Sci U S A 108, 10585–10590 (2011).

14. Jadhav, U. et al. Extensive Recovery of Embryonic Enhancer and Gene Memory Stored in Hypomethylated Enhancer DNA. Mol Cell 74, 542–554.e5 (2019).

15. Zwiggelaar, R. T. et al. LSD1 represses a neonatal/reparative gene program in adult intestinal epithelium. Sci Adv 6, (2020).

16. Ostrop, J. et al. A Semi-automated Organoid Screening Method Demonstrates Epigenetic Control of Intestinal Epithelial Differentiation. Front Cell Dev Biol 8, (2021).

17. Parmar, N. et al. Intestinal-epithelial LSD1 controls goblet cell maturation and effector responses required for gut immunity to bacterial and helminth infection. PLoS Pathog 17, (2021).

18. Shaw, T. N. et al. Tissue-resident macrophages in the intestine are long lived and defined by Tim-4 and CD4 expression. Journal of Experimental Medicine 215, 1507– 1518 (2018).

19. Schneider, C., et al. Tissue-Resident Group 2 Innate Lymphoid Cells Differentiate by Layered Ontogeny and In Situ Perinatal Priming. Immunity 50, 1425–1438.e5 (2019).

20. Mu, Q. et al. Regulation of neonatal IgA production by the maternal microbiota. Proc Natl Acad Sci U S A 118, (2021).

21. Bunker, J. J. & Bendelac, A. IgA Responses to Microbiota. Immunity 49, 211–224 (2018).

22. Rios, D. et al. Antigen sampling by intestinal M cells is the principal pathway initiating mucosal IgA production to commensal enteric bacteria. Mucosal Immunol 9, 907–916 (2016).

23. Knoop, K. A. et al. Microbial antigen encounter during a preweaning interval is critical for tolerance to gut bacteria. Sci Immunol 2, (2017).

24. Kulkarni, D. H. et al. Goblet cell associated antigen passages support the induction and maintenance of oral tolerance. Mucosal Immunol 13, 271–282 (2020).

25. Gerbe, F. et al. Distinct ATOH1 and Neurog3 requirements define tuft cells as a new secretory cell type in the intestinal epithelium. J Cell Biol 192, 767–780 (2011).

26. Von Moltke, J., Ji, M., Liang, H. E. & Locksley, R. M. Tuft-cell-derived IL-25 regulates an intestinal ILC2-epithelial response circuit. Nature 529, 221–225 (2016).

27. El Marjou, F., et al. Tissue-specific and inducible Cre-mediated recombination in the gut epithelium. Genesis 39, 186–193 (2004).

28. Madison, B. B. et al. Cis elements of the villin gene control expression in restricted domains of the vertical (crypt) and horizontal (duodenum, cecum) axes of the intestine. J Biol Chem 277, 33275–33283 (2002).

29. Haber, A. L. et al. A single-cell survey of the small intestinal epithelium. Nature 2017 551:7680 551, 333–339 (2017).

30. Moor, A. E. et al. Spatial Reconstruction of Single Enterocytes Uncovers Broad Zonation along the Intestinal Villus Axis. Cell 175, 1156–1167.e15 (2018).

31. Beumer, J. et al. BMP gradient along the intestinal villus axis controls zonated enterocyte and goblet cell states. Cell Rep 38, (2022).

32. Bahar Halpern, K., et al. Lgr5+ telocytes are a signaling source at the intestinal villus tip. Nat Commun 11, (2020).

33. Beumer, J. et al. Enteroendocrine cells switch hormone expression along the crypt-to-villus BMP signalling gradient. Nat Cell Biol 20, 909–916 (2018).

34. Durand, A. et al. Functional intestinal stem cells after Paneth cell ablation induced by the loss of transcription factor Math1 (Atoh1). Proc Natl Acad Sci U S A 109, 8965– 8970 (2012).

35. Paone, P. & Cani, P. D. Mucus barrier, mucins and gut microbiota: the expected slimy partners? Gut 69, 2232 (2020).

36. Hiippala, K., Kainulainen, V., Kalliomäki, M., Arkkila, P. & Satokari, R. Mucosal prevalence and interactions with the epithelium indicate commensalism of Sutterella spp. Front Microbiol 7, 1706 (2016).

37. Kaiko, G. E. et al. The Colonic Crypt Protects Stem Cells from Microbiota-Derived Metabolites. Cell 165, 1708–1720 (2016).

38. Wu, S. en et al. Microbiota-derived metabolite promotes HDAC3 activity in the gut. Nature 586, 108–112 (2020).

39. Garcia, T. M. et al. Early Life Antibiotics Influence In Vivo and In Vitro Mouse Intestinal Epithelium Maturation and Functioning. Cell Mol Gastroenterol Hepatol 12, 943–981 (2021).

40. Kim, J.-E. et al. Gut microbiota promotes stem cell differentiation through macrophage and mesenchymal niches in early postnatal development. Immunity 55, 2300–2317.e6 (2022).

41. Bain, C. C. et al. Constant replenishment from circulating monocytes maintains the macrophage pool in the intestine of adult mice. Nat Immunol 15, 929–937 (2014).

42. Rivollier, A., He, J., Kole, A., Valatas, V. & Kelsall, B. L. Inflammation switches the differentiation program of Ly6Chi monocytes from antiinflammatory macrophages to inflammatory dendritic cells in the colon. J Exp Med 209, 139–155 (2012).

43. Prise, I. E. et al. CD163 and Tim-4 identify resident intestinal macrophages across sub-tissular regions that are spatially regulated by TGF-β. bioRxiv 2023.08.21.553672 (2023) doi:10.1101/2023.08.21.553672.

44. Serbina, N. V. & Pamer, E. G. Monocyte emigration from bone marrow during bacterial infection requires signals mediated by chemokine receptor CCR2. Nat Immunol 7, 311–317 (2006).

45. Takada, Y. et al. Monocyte Chemoattractant Protein-1 Contributes to Gut Homeostasis and Intestinal Inflammation by Composition of IL-10–Producing Regulatory Macrophage Subset. The Journal of Immunology 184, 2671–2676 (2010).

46. Dick, S. A., et al. Three tissue resident macrophage subsets coexist across organs with conserved origins and life cycles. Sci Immunol 7, (2022).

47. Kotarsky, K. et al. A novel role for constitutively expressed epithelial-derived chemokines as antibacterial peptides in the intestinal mucosa. Mucosal Immunol 3, 40– 48 (2010).

48. Gury-BenAri, M. et al. The Spectrum and Regulatory Landscape of Intestinal Innate Lymphoid Cells Are Shaped by the Microbiome. Cell 166, 1231–1246.e13 (2016).

49. Korbecki, J. et al. The Role of CXCL16 in the Pathogenesis of Cancer and Other Diseases. International Journal of Molecular Sciences 2021, Vol. 22, Page 3490 22, 3490 (2021).

50. Robinette, M. L. et al. Transcriptional programs define molecular characteristics of innate lymphoid cell classes and subsets. Nat Immunol 16, 306–17 (2015).

51. Egolf, S. et al. LSD1 Inhibition Promotes Epithelial Differentiation through Derepression of Fate-Determining Transcription Factors. Cell Rep 28, 1981–1992.e7 (2019).

52. Alenghat, T. et al. Histone deacetylase 3 coordinates commensal-bacteria-dependent intestinal homeostasis. Nature 504, 153–157 (2013).

53. Vaishnava, S. et al. The antibacterial lectin RegIIIgamma promotes the spatial segregation of microbiota and host in the intestine. Science 334, 255–258 (2011).

54. Satoh-Takayama, N. et al. The chemokine receptor CXCR6 controls the functional topography of interleukin-22 producing intestinal innate lymphoid cells. Immunity 41, 776–788 (2014).

55. Pu, Q. et al. Gut Microbiota Regulate Gut–Lung Axis Inflammatory Responses by Mediating ILC2 Compartmental Migration. The Journal of Immunology 207, 257–267 (2021).

56. Meunier, S. et al. Maintenance of Type 2 Response by CXCR6-Deficient ILC2 in Papain-Induced Lung Inflammation. Int J Mol Sci 20, (2019).

57. Li, Y. et al. Kinetics of the accumulation of group 2 innate lymphoid cells in IL-33-induced and IL-25-induced murine models of asthma: a potential role for the chemokine CXCL16. Cell Mol Immunol 16, 75–86 (2019).

58. Shimaoka, T. et al. Cutting Edge: SR-PSOX/CXC Chemokine Ligand 16 Mediates Bacterial Phagocytosis by APCs Through its Chemokine Domain. The Journal of Immunology 171, 1647–1651 (2003).

59. Victor, A. R. et al. IL-18 Drives ILC3 Proliferation and Promotes IL-22 Production via NF-κB. The Journal of Immunology 199, 2333–2342 (2017).

60. Kerenyi, M. A. et al. Histone demethylase Lsd1 represses hematopoietic stem and progenitor cell signatures during blood cell maturation. Elife 2, (2013).

61. Sato, T. & Clevers, H. Primary mouse small intestinal epithelial cell cultures. Methods Mol Biol 945, 319–328 (2013).

62. Moolenbeek, C. & Ruitenberg, E. J. The ‘Swiss roll’: a simple technique for histological studies of the rodent intestine. Lab Anim 15, 57–59 (1981).

63. Webster, H. C., Andrusaite, A. T., Shergold, A. L., Milling, S. W. F. & Perona-Wright, G. Isolation and functional characterisation of lamina propria leukocytes from helminth-infected, murine small intestine. J Immunol Methods 477, (2020).

64. Dobin, A. et al. STAR: ultrafast universal RNA-seq aligner. Bioinformatics 29, 15–21 (2013).

65. Frankish, A. et al. GENCODE 2021. Nucleic Acids Res 49, D916–D923 (2021).

66. Liao, Y., Smyth, G. K. & Shi, W. featureCounts: an efficient general purpose program for assigning sequence reads to genomic features. Bioinformatics 30, 923–930 (2014).

67. GitHub - kevinblighe/EnhancedVolcano: Publication-ready volcano plots with enhanced colouring and labeling. https://github.com/kevinblighe/EnhancedVolcano.

68. Pedregosa FABIANPEDREGOSA, F. et al. Scikit-learn: Machine Learning in Python Gaël Varoquaux Bertrand Thirion Vincent Dubourg Alexandre Passos PEDREGOSA, VAROQUAUX, GRAMFORT ET AL. Matthieu Perrot. Journal of Machine Learning Research 12, 2825–2830 (2011).

69. Yu, G., Wang, L. G., Han, Y. & He, Q. Y. clusterProfiler: an R package for comparing biological themes among gene clusters. OMICS 16, 284–287 (2012).

70. Zheng, G. X. Y. et al. Massively parallel digital transcriptional profiling of single cells. Nat Commun 8, (2017).

71. Wolf, F. A., Angerer, P. & Theis, F. J. SCANPY: large-scale single-cell gene expression data analysis. Genome Biol 19, (2018).

72. Ekiz, H. A., Conley, C. J., Stephens, W. Z. & O’Connell, R. M. CIPR: a web-based R/shiny app and R package to annotate cell clusters in single cell RNA sequencing experiments. BMC Bioinformatics 21, 191 (2020).

73. Magoč, T. & Salzberg, S. L. FLASH: fast length adjustment of short reads to improve genome assemblies. Bioinformatics 27, 2957–2963 (2011).

