## Supplementary Figures for "Postnatal intestinal epithelial maturation by LSD1 controls the small intestinal immune cell composition independently from the microbiota"

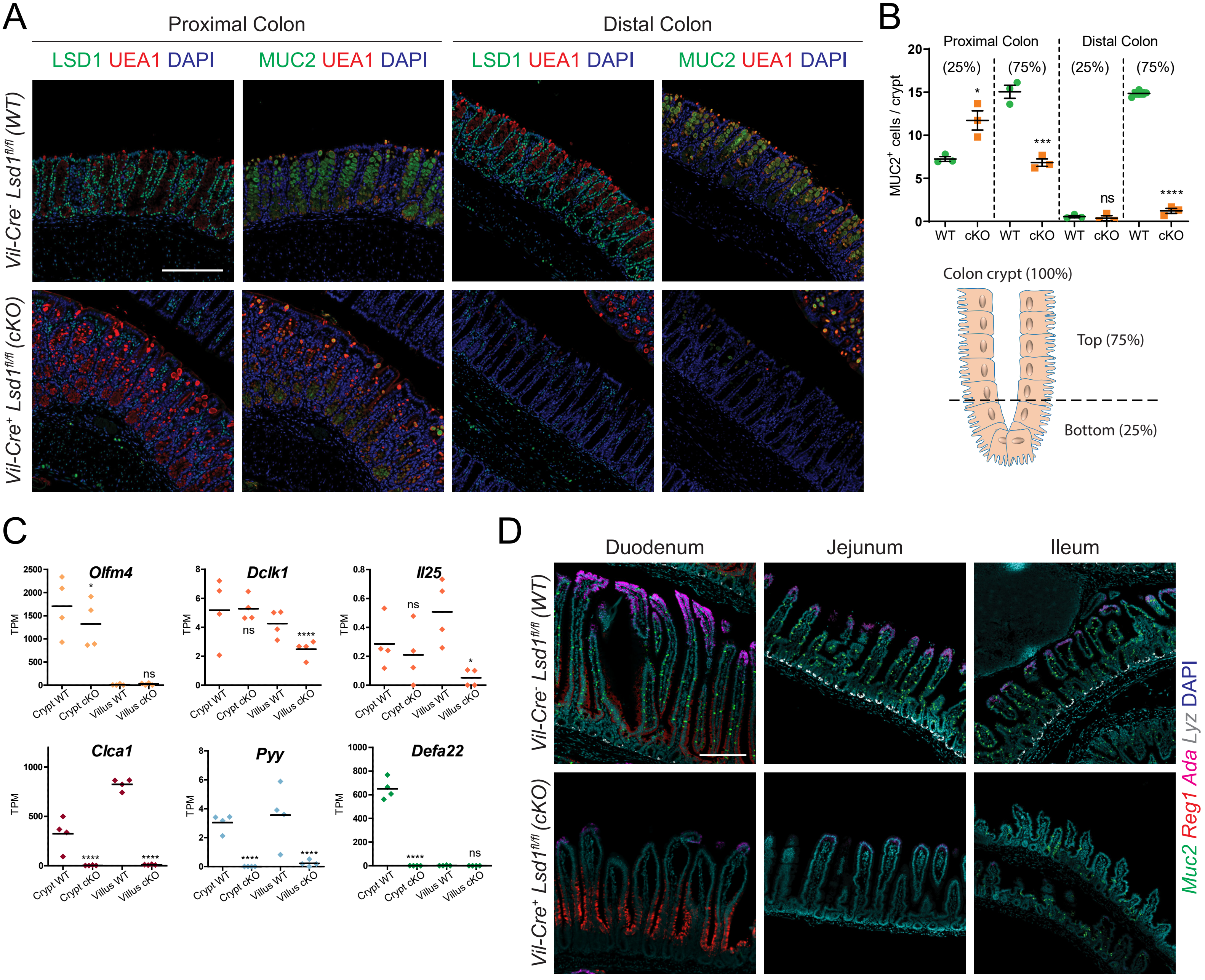


**Fig. S1. LSD1 is required for the postnatal maturation of the intestinal epithelium. (A)** Immunofluorescence staining of paraffin-embedded mouse colon tissue depicting complete loss of intestinal epithelial LSD1, and decreased goblet cells (MUC2+ UEA1+) in Lsd1 cKO mice, particularly in the distal portion. Scale bar: 200µm. **(B)** Quantification of goblet (MUC2+) cells across the colon (derived from Fig. S1A images). In order to depict the accumulation of immature/precursor goblet cells at the bottom of proximal colon crypts, crypts have been segmented according to their bottom 25% and top 75% portions. Data are presented as mean ± SEM in a scatter plot; n = 3 mice/genotype, (Two-tailed unpaired t-test). **(C)** Bulk RNA-seq of crypt and villus fractions derived from WT and cKO 2-month-old mice. Individual graphs show Transcripts per Million (TPM). Data are presented as mean & individual data points; n = 4 mice/genotype, (Benjamini–Hochberg adjusted p-value). **(D)** Representative fluorescence in situ hybridization of key cellular markers for goblet (Muc2, green), Paneth (Lyz, white), villus-base (Reg1, red) and villus-tip (Ada, magenta) enterocytes. Scale bar: 200µm; n = 3 mice/genotype. Cell nuclei in all imaging are counterstained with DAPI.

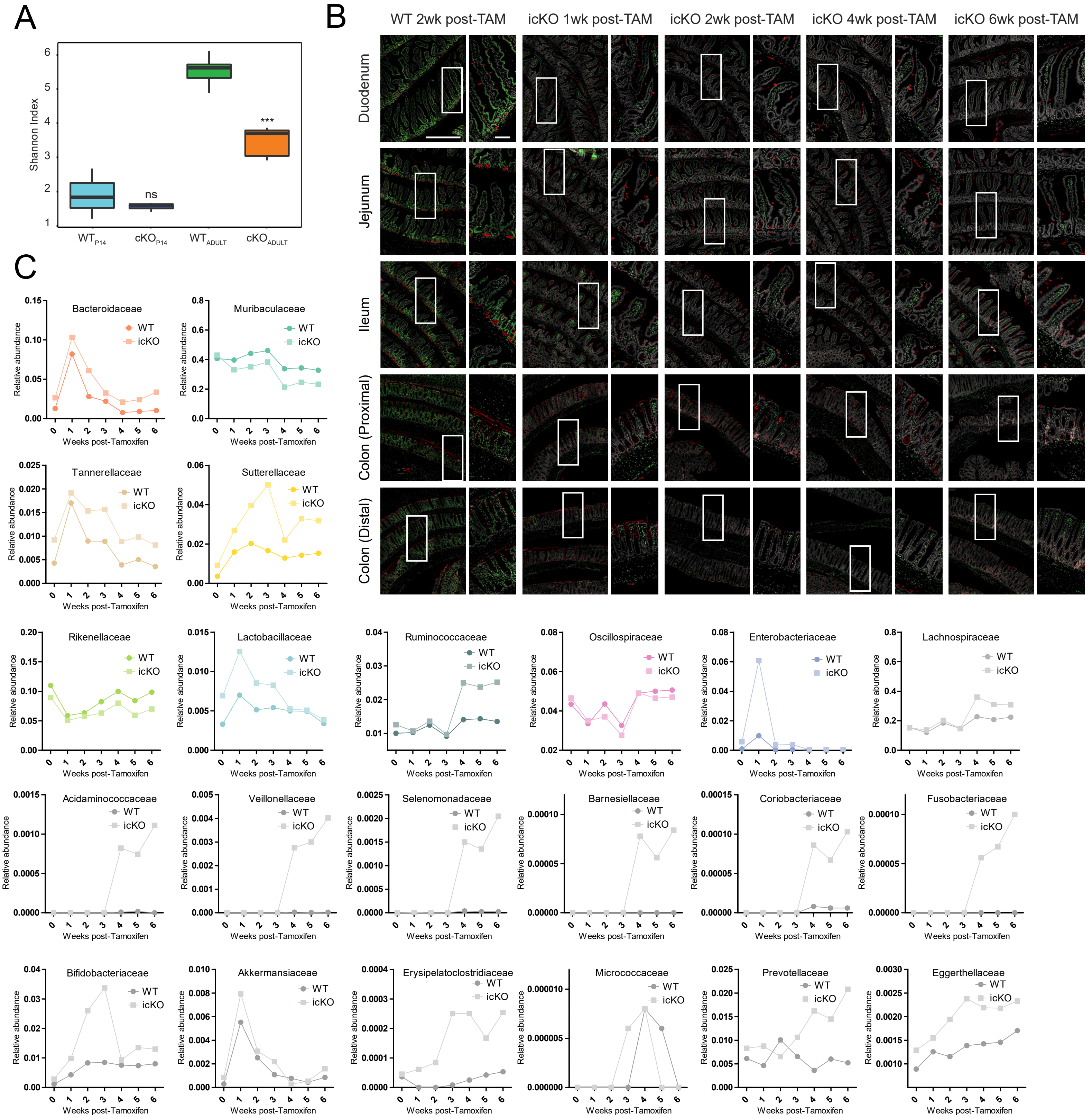


**Fig. S2. Mature intestinal epithelium defines and maintains microbial composition. (A)** Stool microbiota species diversity in adult (2-month-old) and P14 mice (WT and cKO); n = 5 adult mice/genotype & n = 3 P14 mice/genotype (Shannon Diversity Index). **(B)** Immunofluorescence staining of paraffin-embedded mouse intestinal tissue stained for LSD1 and UEA1 after tamoxifen treatment. Scale bar: 500µm, inset scale bar: 100µm; n = 5 mice/genotype from 2 independent experiments. **(C)** Relative abundance representation of selected bacterial families before and after tamoxifen administration in WT and icKO litter and cagemates; n = 6 mice/genotype/timepoint.

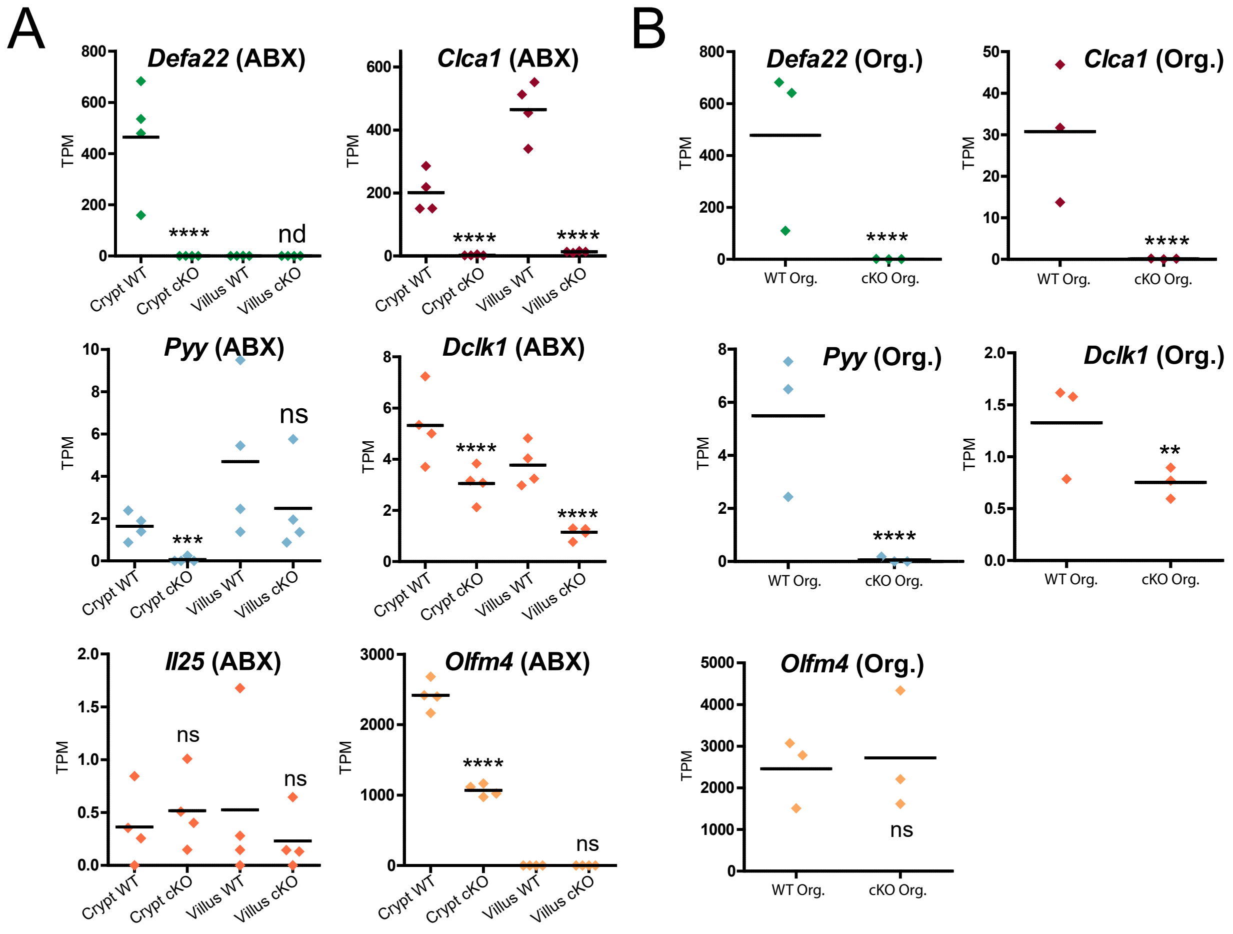


**Fig. S3. LSD1 driven epithelial maturation is independent from the microbiota. (A)** Bulk RNA-seq of crypt and villus fractions derived from WT (ABX) and cKO (ABX) 2-month-old mice. Individual graphs show Transcripts per Million (TPM). Data are presented as mean & individual data points; n = 4 mice/genotype, (Benjamini–Hochberg adjusted p-value). **(B)** Bulk RNASeq of organoids derived from untreated WT and cKO mice. Individual graphs show Transcripts per Million (TPM). Data are presented as mean & individual data points; n = 3 mice/genotype, (Benjamini–Hochberg adjusted p-value).

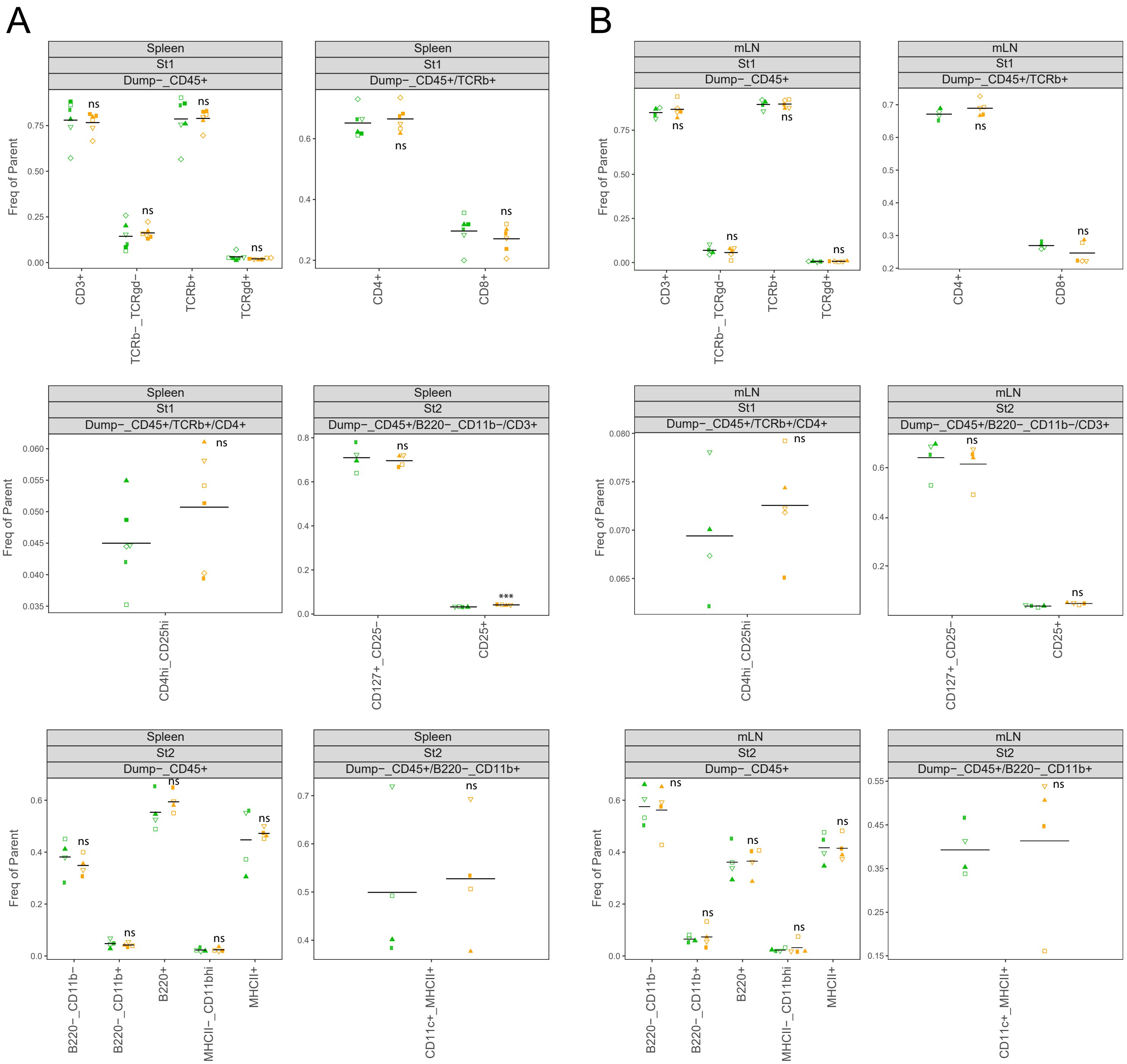


**Fig. S4. LSD1-mediated intestinal epithelial maturation does not control systemic immune cell imbalance in spleen or mesenteric lymph nodes but directs local immune cell populations. (A & B)** Flow cytometry data derived from spleen **(A)** or mLN **(B)** tissues showing frequency of B cells, T cell populations and myeloid cell populations. Green and orange data points correspond to WT and cKO mice respectively; Each data point represents and individual mouse, independent experiments were carried out for each WT and cKO pair, (Two-tailed unpaired t-test). St1 and St2 represent two different staining combinations. Parent gates are shown above each plot. Full gating hierarchy is shown in (S4E). Cells were gated on viable CD45+ cells. Ly6g+ granulocytes were excluded from the analysis. B220 was used as marker for B cells, and CD3, TCRb, TCRgd, CD4, CD8, CD25, CD127 were used to investigate T cell populations including CD4+ and CD8+ T helper cells, CD4hi CD25hi Tregs, and TCRb-TCRgd- “double-negative” T cells. CD11b, CD11c and MHCII were included to investigate myeloid cell populations including CD11c+ MHCII+ dendritic cells.

C

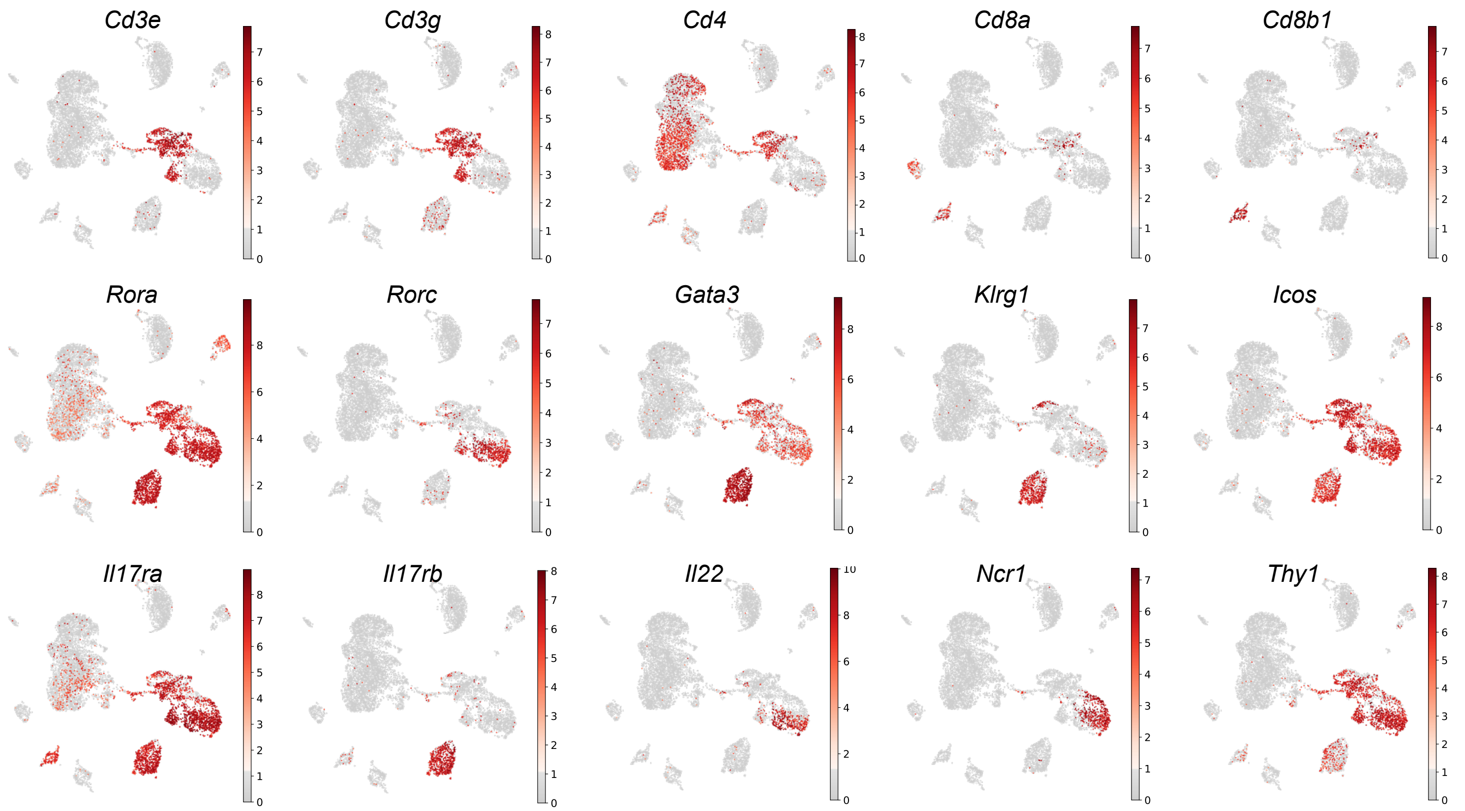


**Fig. S4 (continued). LSD1-mediated intestinal epithelial maturation does not control systemic immune cell imbalance in spleen or mesenteric lymph nodes but directs local immune cell populations. (C)** UMAP of lamina propria CD45+-derived cells depicting individual gene expression across all experimental conditions merged (untreated WT, untreated cKO, ABX WT and ABX cKO). Grey to red heatmap scale shows log(Counts Per Million+1) or log(CPM+1).

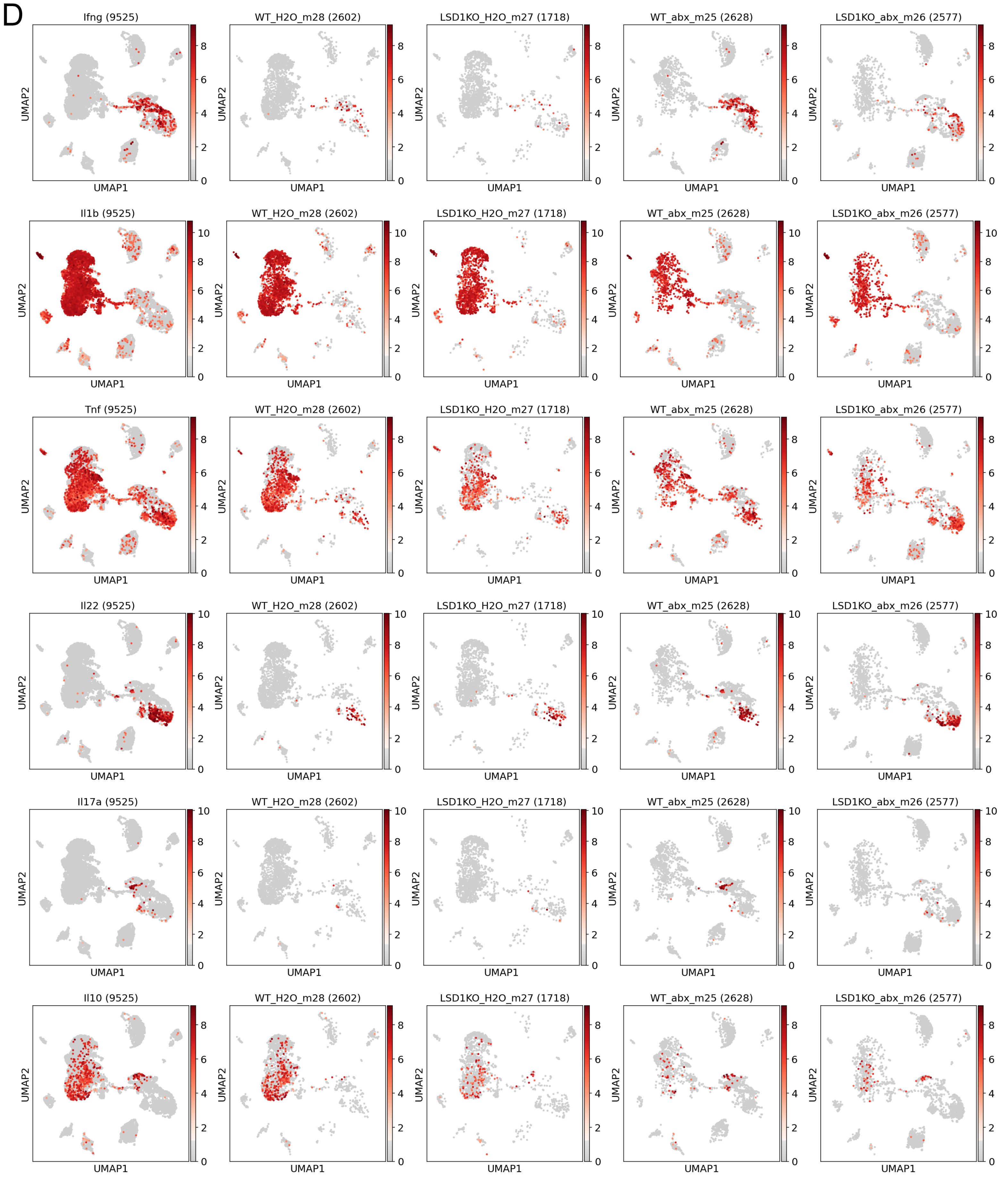


**Fig. S4 (continued). LSD1-mediated intestinal epithelial maturation does not control systemic immune cell imbalance in spleen or mesenteric lymph nodes but directs local immune cell populations. (D)** UMAP of lamina propria CD45+-derived cells showing individual gene expression across all experimental conditions, from left to right: all conditions merged, untreated WT, untreated cKO, ABX WT and ABX cKO. Number in between parentheses represents the number of sequenced cells that passed quality control, (9525) corresponds to all four conditions merged under one UMAP. Grey to red heatmap scale shows log(CPM+1).

E

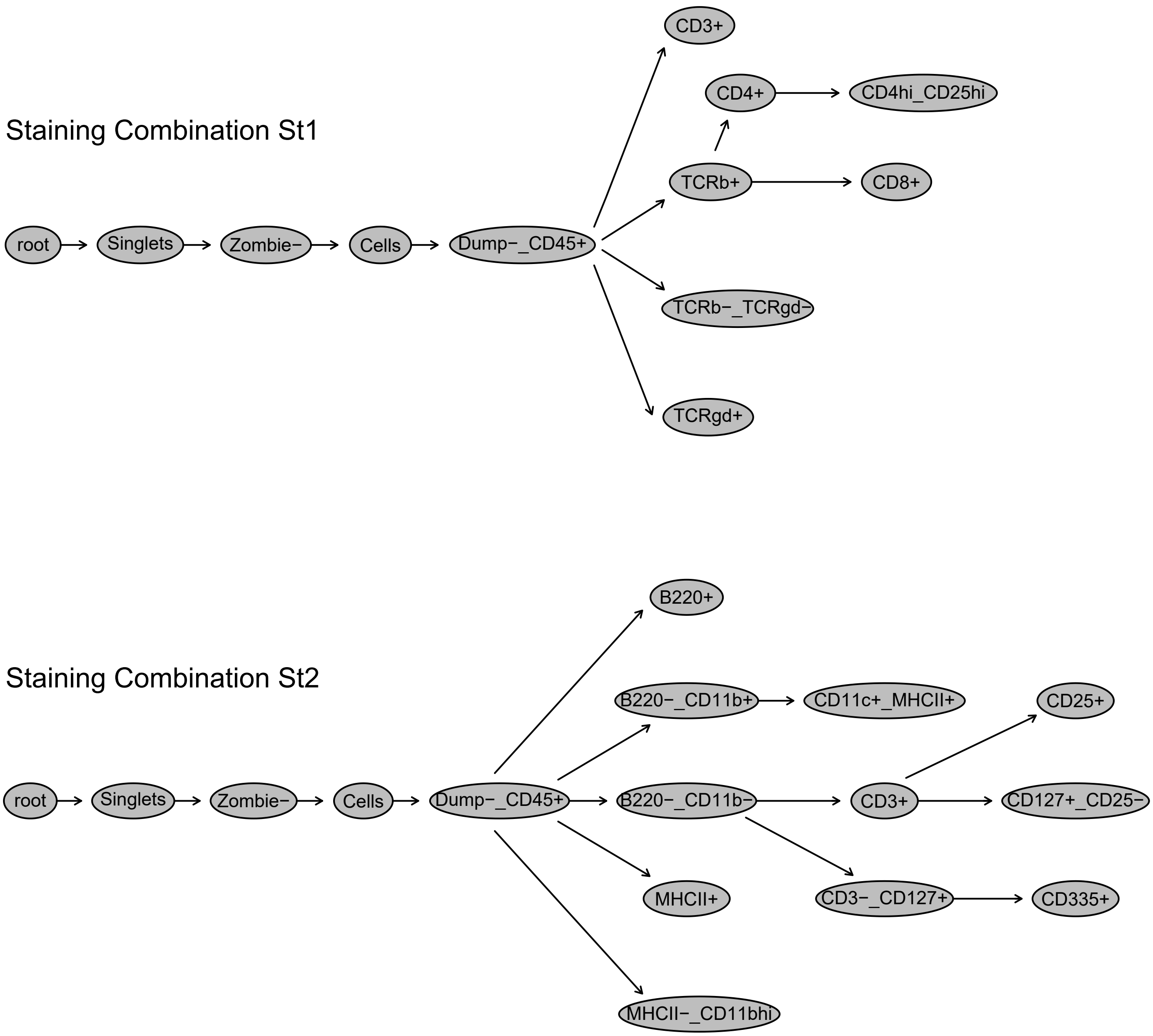


**Fig. S4 (continued). LSD1-mediated intestinal epithelial maturation does not control systemic immune cell imbalance in spleen or mesenteric lymph nodes but directs local immune cell populations. (E)** Complete gating hierarchy for analysis of immune cell populations by staining combinations St1 [CD3-BV605, CD8-BV785, CD25-AF488, TCRgd-PerCp-Cy5.5, TCRb-PE, CD4-APC, CD45-APC-Fire, Dump (CD326, CD19, CD11b, Ly6g, Ter119)-PE-Cy7] and St2 [CD335-BV421, CD3-BV605, CD127-BV711, CD25-BV785, MHCII-AF488, CD11c-PerCp-Cy5.5, B220-PE, CD11b-AF647, CD45-APC-Fire, Dump(CD326, Ly6g, Ter119)-PE-Cy7].

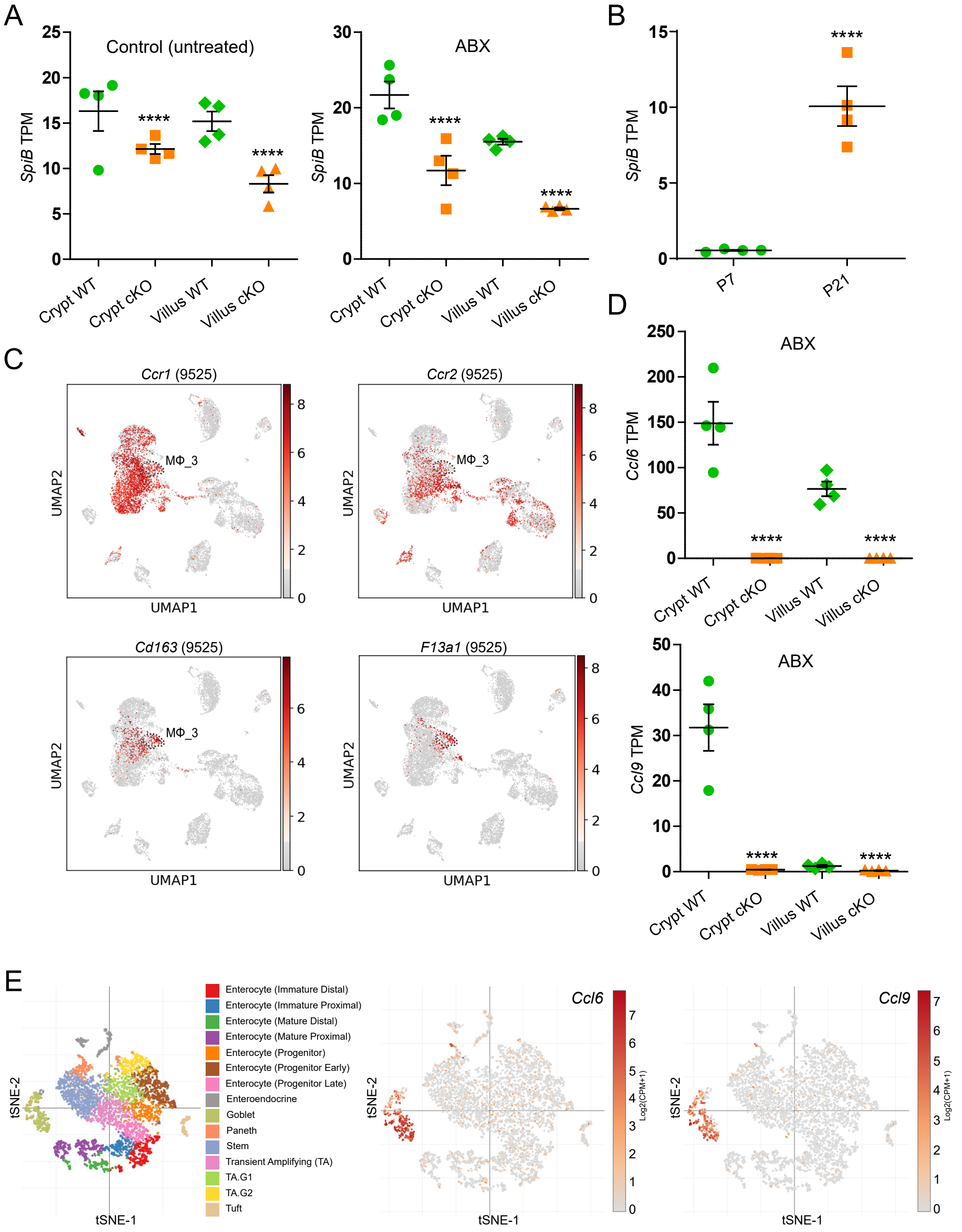


**Fig. S5. LSD1-mediated intestinal epithelial maturation controls intestinal plasma cell and macrophage homeostasis. (A)** Bulk RNA-seq of crypt and villus fractions derived from untreated and ABX-treated 2-month-old mice. Individual graphs show Transcripts per Million (TPM). Data are presented as mean ± SEM; n = 4 mice/genotype/condition, (Benjamini–Hochberg adjusted p-value). **(B)** Bulk RNA-seq of intestinal epithelium derived from WT mice at postnatal day 7 (P7) vs P2110. Individual graphs show Transcripts per Million (TPM). Data are presented as mean ± SEM; n = 4 mice/timepoint, (Benjamini–Hochberg adjusted p-value). **(C)** UMAP of lamina propria CD45+-derived cells showing Ccr1, Ccr2, Cd163 and F13a1 gene expression across all experimental conditions. Number in between parentheses represents the number of sequenced cells that passed quality control, (9525) corresponds to all four conditions merged under one UMAP. Grey to red heatmap scale shows log(CPM+1). **(D)** Bulk RNA-seq of crypt and villus fractions derived from WT and cKO ABX-treated 2-month-old mice. Individual graphs show Transcripts per Million (TPM). Data are presented as mean ± SEM; n = 4 mice/genotype, (Benjamini–Hochberg adjusted p-value). **(E)** UMAP plot of cell types as determined with scRNA-seq from cells derived from the small intestinal epithelium of mice. Each cell type class is represented as a cluster of points in a unique color (left panel). UMAP showing Ccl6, and Ccl9 gene expression across cell types (middle and right panels). Grey to red heatmap scale shows log(CPM+1). Data derived from[29].

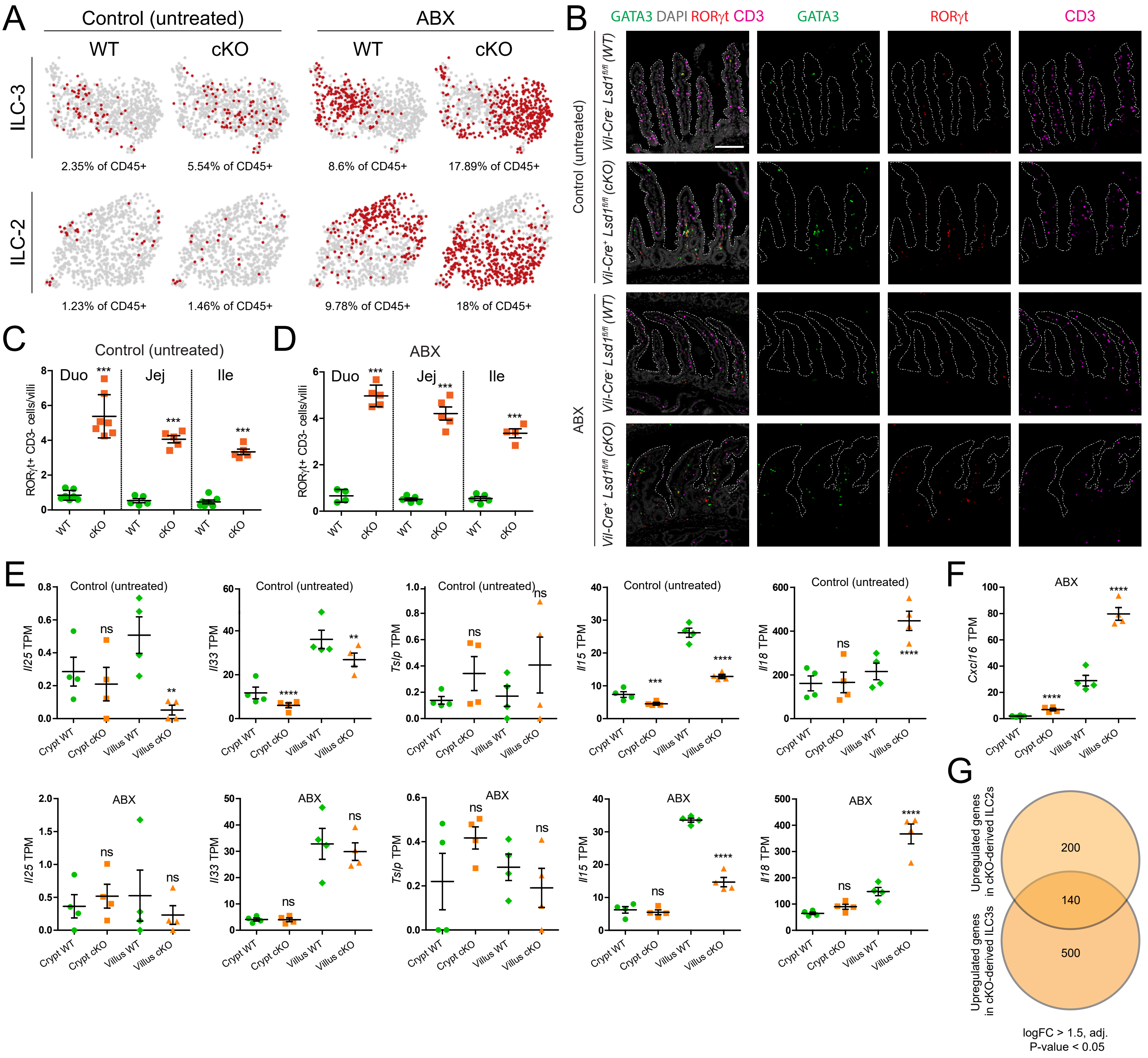


**Fig. S6. LSD1-mediated intestinal epithelial maturation is required for the establishment and maintenance of ILC2s and ILC3s in a microbiota-independent manner. (A)** UMAP clustering of lamina propria ILC2s and ILC3s across all experimental conditions. Red dots correspond to the number of cells detected under each condition. Grey dots represent the sum of all detected ILC2 or ILC3 cells across conditions. **(B)** Split channel immunofluorescence of mouse duodenums derived from Fig. 6A. ILC2s are defined as CD3- RORγt- GATA3+ while ILC3s are CD3- RORγt+. CD3 (magenta), RORγt (red), GATA3 (green) and nuclei are counterstained with DAPI (grey). Villus structure is delimited by a discontinuous white line; Scale bar: 200µm. **(C)** Quantification of ILC3s in the lamina propria of untreated mice across the small intestine. Data are presented as mean ± SEM; n = 7 mice/genotype (duodenum) and 5 mice/genotype (jejunum & ileum) from 2 independent experiments, (Two-tailed unpaired t-test). **(D)** Quantification of ILC3s in the lamina propria of ABX-treated mice across the small intestine. Data are presented as mean ± SEM; n = 5 mice/genotype from 2 independent experiments, (Two-tailed unpaired t-test). **(E & F)** Bulk RNA-seq of crypt and villus fractions derived from WT and cKO untreated or ABX-treated 2-month-old mice. Individual graphs show Transcripts per Million (TPM). Data are presented as mean ± SEM; n = 4 mice/genotype, (Benjamini–Hochberg adjusted p-value). **(G)** Venn diagram showing overlapping upregulated genes found in both ILC2s and ILC3s derived from ABX treated mice.

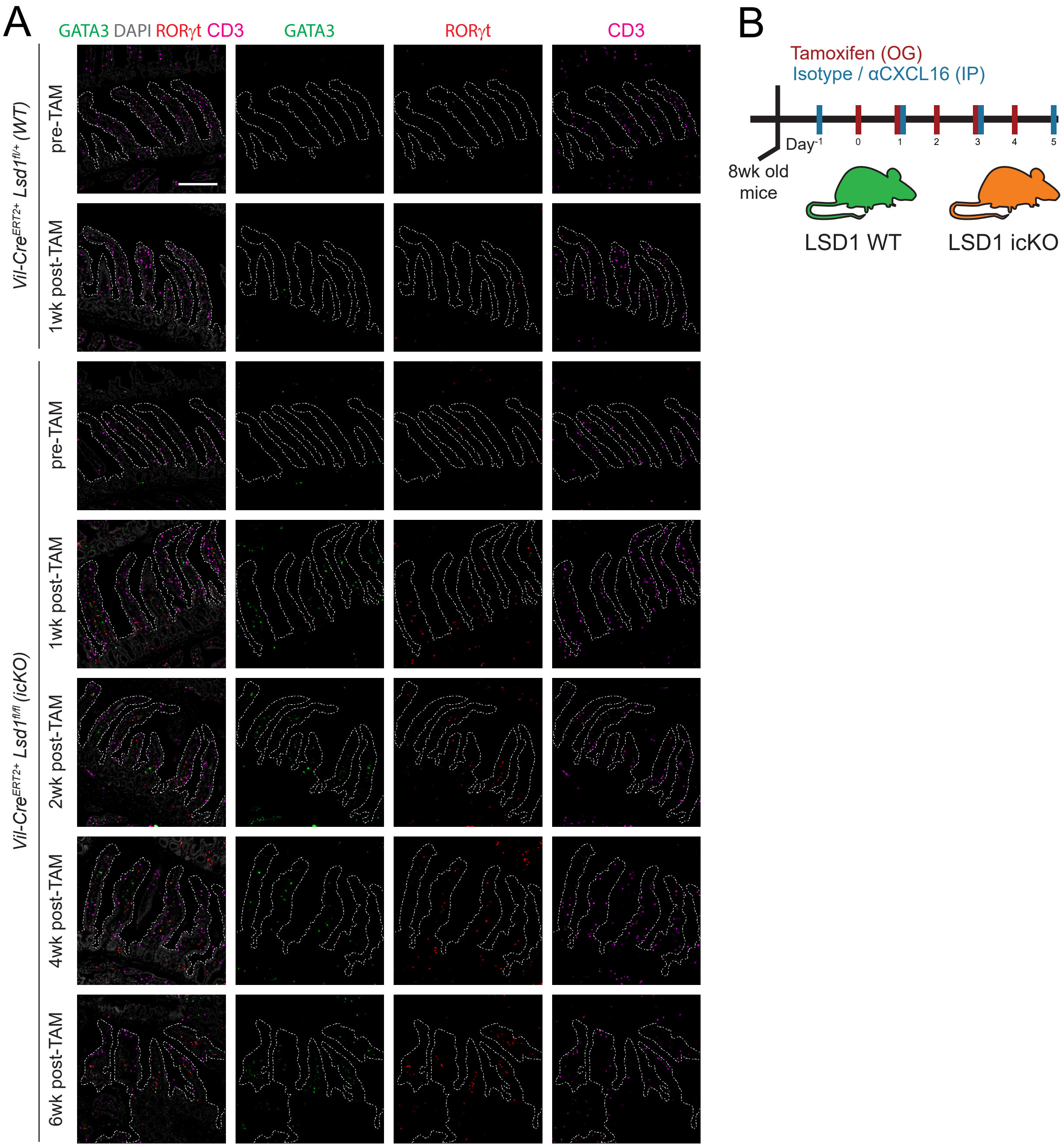


**Fig. S7. Rapid induction of CXCL16-dependent ILC2s and CXCL16-independent ILC3s upon epithelial-intrinsic Lsd1 deletion in adult mice. (A)** Split channel immunofluorescence of mouse duodenums derived from Fig. 7A. ILC2s are defined as CD3- RORγt- GATA3+ while ILC3s are CD3- RORγt+. CD3 (magenta), RORγt (red), GATA3 (green) and nuclei are counterstained with DAPI (grey). Villus structure is delimited by a discontinuous white line; Scale bar: 200μm. **(B)** Treatment schematics for the CXCR6-CXCL16 signaling blockade experiment (α CXCL16). See materials and methods for a detailed protocol.
